# Predictive Coding of Reward in the Hippocampus

**DOI:** 10.1101/2024.09.03.611040

**Authors:** Mohammad Yaghoubi, Andres Nieto-Posadas, Coralie-Anne Mosser, Thomas Gisiger, Émmanuel Wilson, Sylvain Williams, Mark P. Brandon

## Abstract

A fundamental objective of the brain is to anticipate future outcomes^1–7^. This process requires learning the states of the world as well as the transitional relationships between those states. The hippocampal cognitive map is believed to be one such internal model^8^. However, evidence for predictive coding^9–12^ and reward sensitivity^13–19^ in the hippocampal neuronal representation suggests that its role extends beyond purely spatial representation. In fact, it raises the question of what kind of spatial representation is most useful for learning and maximizing future rewards? Here, we track the evolution of reward representation over weeks as mice learn to solve a cognitively demanding reward-based task. Our findings reveal a highly organized restructuring of hippocampal reward representations during the learning process. Specifically, we found multiple lines of evidence, both at the population and single-cell levels, that hippocampal representation becomes predictive of reward over weeks. Namely, both population-level information about reward and the percentage of reward-tuned neurons decrease over time. At the same time, the representation of the animals’ choice and reward approach period (the period between choice and reward) increased over time. By tracking individual reward cells across sessions, we found that neurons initially tuned for reward shifted their tuning towards choice and reward approach periods, indicating that reward cells backpropagate their tuning to anticipate reward with experience. These findings underscore the dynamic nature of hippocampal representations, highlighting their critical role in learning through the prediction of future outcomes.

## Introduction

The hippocampus represents a mixture of spatial (such as place^20^, Landmarks^21^, etc.) and non-spatial (such as time^22^, sound frequency^23^, odor^24^, lap number^25^, reward^13,18,26,27^, etc.) environmental features. The collective encoding of environmental features and their relationships, known as a cognitive or hippocampal map^8^, is believed to support spatial navigation and memory-related behaviors — cognitive processes essential for an animal’s survival. From an evolutionary standpoint, an animal’s survival depends on using these cognitive abilities to efficiently learn and remember rewarding experiences, such as navigating to home, safety, and food. Consequently, the hippocampal representation of the environment is expected to undergo significant changes once an animal has learned how to navigate to rewarding locations or has mapped environmental features associated with rewards^26^.

Previous work has examined the modulation of hippocampal representation at various time points during reward learning, such as reward approach, onset, location, and history^13^. For reward approach, it has been shown that running towards a known goal location induces place-specific firing along paths to goals that are distinct from firing along the same paths during random foraging in the same environment^17^. Place fields also have been observed to cluster near reward locations, resulting in an overrepresentation of those locations by the neural population^14–16^. At the reward arrival, a distinct population of hippocampal neurons has been shown to consistently encode for reward delivery, despite changes in location or context. This suggests that a hippocampal reward signal can be dissociated from place firing^18^. It has also been shown that the hippocampus encodes reward history. In particular, depending on the outcome of the reward, the hippocampal cells modulate their firing activity after the probabilistic delivery of the reward and after leaving the reward site^19^. These studies have examined changes in hippocampal neuronal representation before and after learning reward location, but how these dynamics emerge with extended experience (across days, weeks, or months) remains unknown.

The hippocampus has been shown to support predictive models in various species^9,11,28–36^. Learning to predict rewards is crucial for an animal’s survival. Therefore, we propose that the reorganization of hippocampal representations, in particular reward representation, during learning a reward-based task will occur to enhance reward prediction. We examine this hypothesis by tracking the evolution of the hippocampal representation across weeks as mice perform a reward-based delayed nonmatching-to-location task.

## Results

### Calcium imaging of CA1 neurons across an extended learning task

We used a head-mounted miniscope^37,38^ to perform calcium imaging of the dorsal CA1 hippocampus in eight mice (Fig. 1a). Calcium imaging data were preprocessed with open-source code called NoRMCorre to correct for the motion artifacts^39^, and CNMFe to segment cells and extract calcium transients^40,41^ (Fig. 1b-c, Extended Data Fig. 1). Calcium transients were deconvolved by fitting a generative autoregressive model of calcium dynamics to estimate spiking probabilities of individual cells^42^ (Fig. 1c). Our recording apparatus enabled us to record 504 ± 101 (mean ± std) neurons across sessions and mice (Fig. 1d). We used an automated touchscreen recording box^43,44^, a 20 x 18 cm chamber that consists of a touchscreen in front, a reward port in the back, and an Infrared (IR) camera on top to record behavior (Fig. 1e). Mice were trained to perform a delayed nonmatching-to-location task. After initiation of the task, a sample is presented on one of the screens (either left or right). Following nose poke to the sample, the delay starts, and two white squares are displayed after the delay. Mice must choose the non-matching square to receive the reward located on the back wall of the touchscreen chamber (Fig. 1e-f). We increased the difficulty of the task by increasing the delay period between sample and choice initiation as the mouse reached a high-performance criterion for two reasons. First, in most experimental studies, time and learning (measured by the mouse’s performance) are strongly correlated, making it challenging to separate the effects of time (session) and performance on changes in neuronal activity. Our approach addresses this issue, allowing us to study the effects of time and mouse performance on the neuronal dynamics of hippocampal cells separately. Second, progressively making the task more difficult ensures that the mouse continually learns, engaging the neuronal circuits involved in learning throughout the entire recording period. Previous studies demonstrate that hippocampal lesions impair mouse behavior in learning Trial-unique, delayed nonmatching-to-location, with longer delays exacerbating the deficit45. The learning curve in Fig. 1g shows mouse performance (proportion of correct trials) vs time (session) (see also Extended Data Fig. 2).

**Fig. 1.**
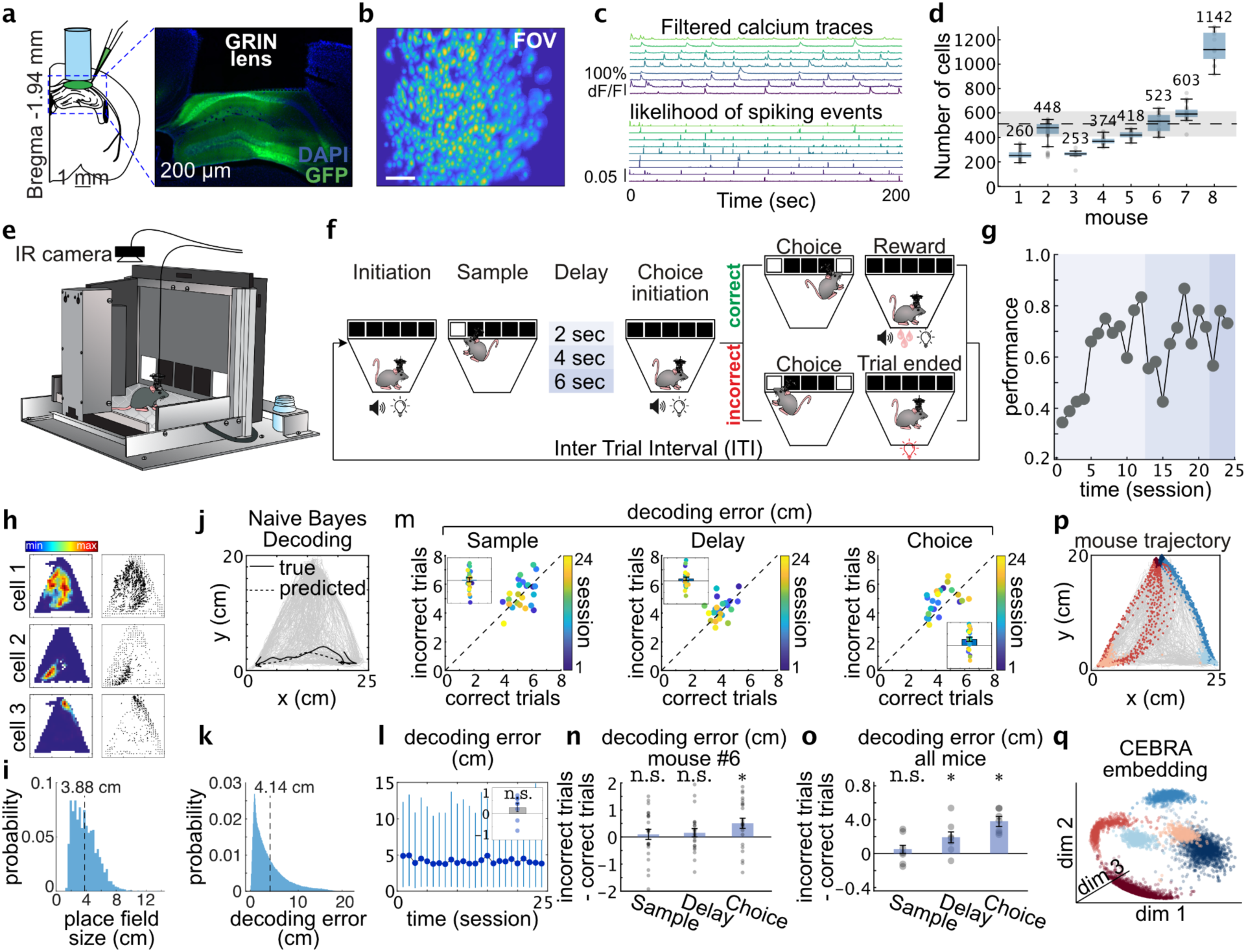
Imaging CA1 neuronal activity in mice while performing a reward-based task. **a,** Schematic of the surgical strategy. **b,** Field of view (FOV) of CA1 of a given session showing the identified cells (FOVs of all mice are presented in Extended Data Fig. 1). **c,** Extracted calcium traces of 9 representative cells (top rows) and their corresponding deconvolved traces demonstrate the inferred likelihood of spiking events (bottom rows). **d,** The number of identified cells across 188 sessions and eight mice (The number of recorded sessions for our eight mice is as follows: 33, 53, 17, 18, 7, 24, 23, 13). Each point represents one recording session. The dashed gray line and the shade around it represent the mean ± std (504 ± 101) of the number of cells. **e,** Schematic of the touchscreen chamber. **f,** Schematic of the task. Briefly, after initiation, in the sample phase, a white square is presented in one of two positions on the touchscreen - either the most left or the most right. A nose poke to this square starts a delay period (increasing from two to six seconds as the mice learn the task). Following this delay, two white squares (most left and most right) are displayed, and the mouse must choose the nonmatching square to receive the reward, located on the back wall of the touchscreen chamber. **g,** Learning curve demonstrating mouse performance (proportion of correct trials in each session) vs time (session) for a representative mouse with 24 sessions. (see Extended Data Fig. 2). **h,** Place fields of three representative cells (left column) and their corresponding vector fields (right column). **i,** Distribution of the size of the place fields of identified place cells is plotted. The average place field size is ∼3.88 cm (see Extended Data Fig. 3). **j,** Naive Bayes (NB) position decoding shows an accurate position decoding from raw calcium traces. All registered cells in each session are used to train the NB model. **k,** Distribution of decoding error across frames of 24 sessions. Mouse #6 shows a heavy tail distribution. The dashed line shows the mean decoding error: ∼4.14 cm. **l,** Decoding error (averaged within each session) over time for mouse #6 was calculated. Each point represents the average decoding error across all frames in a session, with error bars indicating the 95% confidence interval of the lower and upper bounds of the decoding error distribution. The inset shows the correlation of decoding error with time (session) across mice. We do not observe a significant trend across mice. **m,** Comparing decoding error of position for correct and incorrect trials for three phases of Sample, Delay, and Choice. Note that from sample to choice, the points are shifted above the diagonal, indicating a higher decoding error for incorrect trials. **n,** The decoding analysis suggests that incorrect trials are associated with higher decoding errors as we approach the choice. Each point represents one session of mouse #6 (no. of sessions = 24). P-values of Wilcoxon signed-rank test against zero: P-value (sample) = 0.5633, P-value (delay) = 0.3944, P-value (choice) = 0.0163. **o,** Same analysis as (l) across eight mice. Each point represents a mouse. P-values of Wilcoxon signed-rank test against zero: P-value (sample) = 0.5781, P-value (delay) = 0.0469, P-value (choice) = 0.0156. **p,** The salient moments of the task are color-coded (the moments of touching screen, reward approach, and reward consumption). Each point is a frame. **q,** CEBRA embedding neuronal traces reveal distinct neuronal state spaces for task phases (color coding is the same as panel p). This latent representation is a core concept for some of our later analyses.

Visualization of the calcium rate map of hippocampal cells revealed that individual cells encode the animal’s location within the touchscreen chamber with very small place fields, 3.9 ± 0.1 cm in diameter (Fig. 1h-i, Extended Data Fig. 3). This is particularly interesting because previous studies recorded in open fields typically report hippocampal place fields to be around 20 cm in size^46–50^. Using an information theory-based shuffle-control procedure; we found that 48 ± 2% of the cells are identified as place cells. Moreover, the recruitment of place cells shows a fairly consistent trend across time (Extended Data Fig. 4). The tuning properties of place cells showed a strong modulation by behavior (Extended Data Fig. 5). We trained a Bayesian decoder to estimate the animal’s position and extract the spatial information content from the neuronal population^51^ (Fig. 1j). The population spatial decoding results show that in smaller environments, such as our touchscreen chamber, the hippocampus encodes space with high precision, achieving a mean decoding error of 4.14 cm (median ∼2.96 cm) (Fig. 1k). Previous studies using state-of-the-art decoding techniques have reported a decoding error of around 10 cm^52,53^. Plotting within-session shows a relatively consistent average decoding error across time (sessions) (Fig. 1l). Furthermore, we compared the decoding errors in correct and incorrect trials in different phases of the task, namely sample, delay, and choice (Fig. 1m). We observed that correct trials have a significantly lower decoding error than incorrect trials (Fig. 1m-n). The difference becomes increasingly pronounced as the animal approaches the choice moment (Fig. 1n). This result is consistent across mice (Fig. 1o). Additionally, and consistent with prior studies^14,54^, our analysis of spatial coding within the touchscreen chamber revealed an over-representation of the reward area once the animal has learned the reward location (Extended Data Fig. 6). Finally, our analysis shows contextual aspects of the task such as correctness of the trials or trial types are encoded in the neuronal population independent of space (Extended Data Fig. 7).

Projecting neuronal traces in a latent space enhances our ability to capture a richer representation, revealing task features encoded more distinctly^55,56^. To extract the low-dimensional structure of the hippocampal representation, we used CEBRA^57^, a recent state-of-the-art technique for embedding large neuronal population activity into a consistent low-dimensional space. The embedding successfully learns the hippocampal internal structure and reveals a distinct state space for different phases of the task, such as screen, reward approach, and reward moments (Fig. 1p-q). We used this embedding space to quantify the information content of the representation of pre-reward cues and the reward itself.

### Reward encoding decreases across experience

We investigated the dynamics of reward representation in the hippocampus as mice learn to solve the nonmatching-to-location task. The learning period varied, taking a few weeks depending on each mouse’s learning rate (Extended Data Fig. 2). To quantify the reward encoding signal across sessions, we conducted analyses at both the population and the single-cell levels. For the population-level analysis, we quantified the information content of the reward representation by using an information theory-based approach using the CEBRA extracted latent space. This systematic framework allowed us to compare reward representation across sessions and observe changes with respect to time and mouse performance. At the single-cell level, we used a shuffle-control approach to identify reward cells in each session and track the evolution of the percentage of reward cells as a function of time and mouse performance. Both analyses suggest that the representation of reward decreases over time and is not correlated with mouse performance.

Before detailing these analyses, we first examined the simple features of raw data during reward consumption. Raw calcium data show dedicated subpopulations of neurons are responsive to reward (Fig. 2a). Reward cells are sorted at the top of the raster plot (Fig. 2a). Notably, the neuronal representation of reward in the hippocampus is modulated by behavior, with distinct subpopulations encoding reward depending on whether the mouse approached the reward port (located at the back of the chamber) after choosing the left or right screen. The sorted calcium traces show that reward neurons do not necessarily align with the onset of the reward^18^ but reliably form a sequence spanning the entire duration of reward consumption (Fig. 2a, Extended Data Fig. 8). We examined the structure of neuronal responses during reward consumption because our experimental setup allowed us to detect the start and end of reward consumption, which lasted around 3-4 seconds. The sequence during reward consumption was reliably observed across trials (Fig. 2a). The average calcium traces across all cells show an increase at reward (Fig. 2b).

**Fig. 2.**
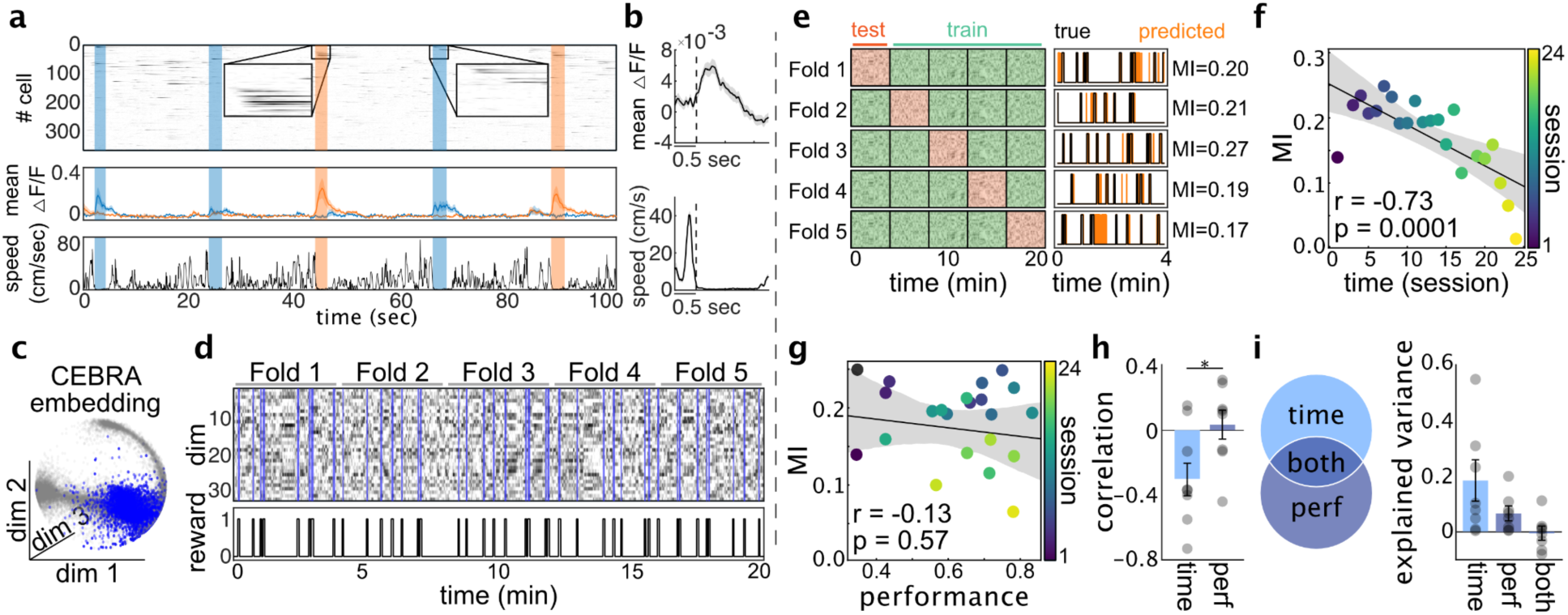
Dynamics of population-level reward coding during learning. **a,** Top panel: Raw calcium traces of 350 recorded cells. The blue and orange bands indicate moments of reward consumption. Orange signifies that the mouse approached the reward port from the left, while blue suggests an approach from the right. The first 45 cells are identified as reward cells and are sorted based on their peak activity during reward consumption. We observe that left and right approaches to the reward engage distinct subpopulations of neurons: cells 1 to 20 are active during right approaches, while cells 21 to 45 are active during left approaches. Notably, each subpopulation forms a neuronal sequence during the reward consumption period, consistently repeated across trials. Middle panel: The average calcium activity for each left and right reward cell subpopulation is plotted. Bottom panel: The speed profile of the mouse is displayed, showing that, as expected, the mouse remains immobile during the reward consumption period. **b,** Top panel: Average calcium activity across all cells at the onset of reward (mean ± SEM, calculated by averaging activity across 35 instances of reward consumption - this session had 35 correct trials). The dashed line shows the onset of the reward consumption. Bottom panel: Average speed profile of mouse (mean ± SEM, averaged across 35 instances of reward consumption). Panels **c**-**e** illustrate our framework for measuring the reward information content encoded in the neuronal population: **c,** An example of neuronal latent representation using CEBRA embedding is presented. each point corresponds to the embedding of neuronal activity with CEBRA from a single frame.; blue points represent frames during reward consumption, while grey points represent all other frames. **d,** The top panel displays the latent representation of the neuronal data obtained using CEBRA embedding, unfolded over time. For all our latent space analyses, the dimensionality of the latent space is 32. The bottom panel shows a binary time series indicating reward moments, with reward frames as 1 and all other frames as 0. **e,** A 5-fold cross-validation is applied to the latent representation of neuronal activity. For each fold, a linear classification model is trained on the other four folds to predict the reward in the held-out data. The Mutual Information (MI) is then calculated between the actual and predicted reward traces. The average MI across the five folds represents the information content of the reward representation for each recording session. **f,** MI vs time (session) is plotted. The negative correlation of MI and time reveals that the information content of reward representation decreases over time. The correlation coefficient = −0.73 (P-value = 0.0001). **g,** MI vs mouse performance is plotted. Mouse performance shows only a weak correlation with MI. The correlation coefficient = −0.13 (P-value = 0.57). **h,** Correlation analysis across all mice (n = 8) indicates that reward MI in hippocampal neurons negatively correlates with time, showing a reduction over time during learning and remaining unaffected by performance. Wilcoxon signed-rank test P-value = 0.0234. **i,** Linear modeling of the reward MI as a function of time and mouse performance reveals that time is the dominant variable that explains most of the variance in the evolution of reward MI. The variance explained by time = 0.17 ± 0.07; the variance explained by performance = 0.07 ± 0.02.

The speed profile confirms that mice remain stationary while receiving the reward, ensuring that the neuronal activity at reward is not driven by mouse behavior (Fig. 2b).

Using CEBRA^57^, we projected our deconvolved calcium traces into a 32-dimensional latent space. Visualization of the first three PCA components of latent space traces reveals a distinct state space for reward representation (Fig. 2c). To quantify the information content of reward representation, we utilized a 5-fold cross-validation approach to decode the reward moments from latent representations. The cross-fold-average mutual information (MI) between the decoded reward traces and the actual reward traces was regarded as the information content of reward representation for each session (Fig. 2d-e). Correlating reward information with time and mouse performance indicates a negative correlation with time (Fig. 2f) and a weaker correlation with mouse performance (Fig. 2g) – suggesting a declining trend in the amount of reward information content over time. The same analysis across all mice shows the same trend (Fig. 2h). Moreover, a linear model assessing reward information content in relation to time and mouse performance indicates that a significant portion of the variance in the dynamics of reward information content is attributed to time rather than mouse performance (Fig. 2i).

To complement this finding, we performed a parallel analysis and examined the evolution of reward encoding at the single-cell level. Reward cells were identified using a shuffling procedure with the threshold of the 99th percentile (Fig. 3a), resulting 8.5 ± 1.5% of the cells (varies across mice and sessions) being identified as reward cells (Fig. 3b). The single-cell tuning curves for reward cells depicted in Fig. 3c reveals two key features: First, the response of individual reward cells is modulated by behavior – meaning that individual cell response to the reward depends on whether the mouse approached the reward port (located at the back of the chamber) after choosing the left screen or right screen. Secondly, different reward cells exhibit tuning to various moments of reward consumption rather than strictly being aligned with reward onset. Furthermore, reward cells showed a significantly higher mean activity (averaged deconvolved traces) during the task than during Inter Trial Intervals (ITI), indicating the task engaged neuronal responses (Fig. 3d, Extended Data Fig. 9). Most importantly, and similar to the results from population-level analysis, correlating the proportion of reward cells with time (session) and mouse performance indicates a negative correlation with time (Fig. 3e) – meaning reduction in recruitment of reward cells across time – and only a weak correlation with mouse performance (Fig. 3f). These results are consistent across mice (Fig. 3g). A linear model assessing the proportion of reward cells in relation to both time and mouse performance indicates that a significant portion of the variance in the dynamics of the proportion of reward cells is attributed to time (session) rather than mouse performance (Fig. 3h). Both population level and single-cell level analysis reveal that the reward representation decreases over time. The observed structured and gradual reorganization of reward representation in the hippocampus prompted us to extend our analysis to examine the dynamics of hippocampal representation for other salient features of the task, such as the pre-reward epochs, over time.

**Fig. 3.**
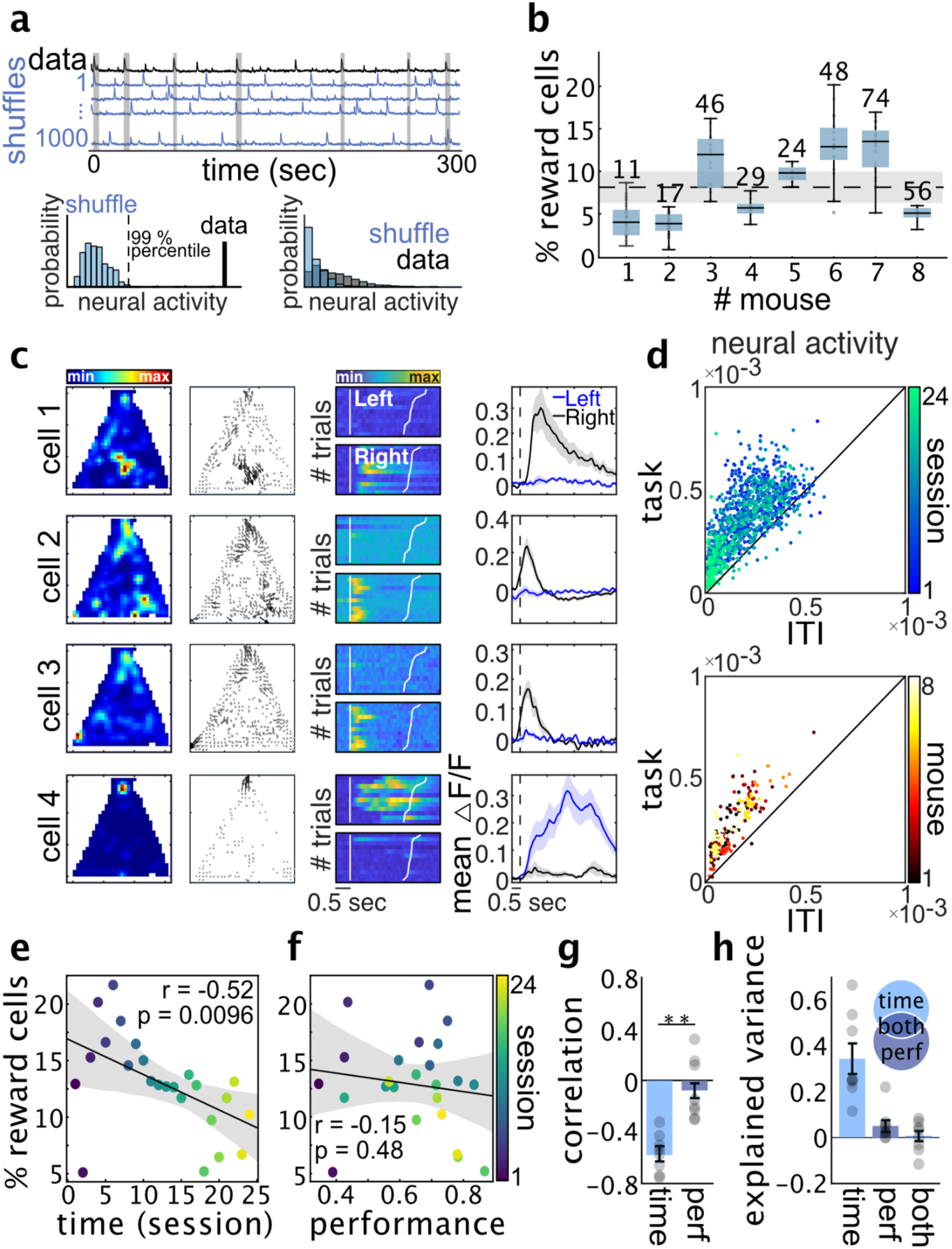
Dynamics of reward cell recruitment during learning. **a,** The shuffle control procedure is employed to identify reward cells. It involves calculating the neuronal activity during reward consumption (averaging the deconvolved trace) for each neuron and comparing it with the distribution of neuronal activity from 1000 random circular shuffled traces. A neuron is classified as a reward cell if its neuronal activity surpasses the 99th percentile of the distribution of shuffled neuronal activity (bottom left panel). The bottom right panel shows the distribution of neuronal activity for all reward cells (black) and compares them with their corresponding shuffled ones (blue) (n = 6495). **b,** Percentage of identified reward cells across mice. Each point represents one session, and the numbers above each box show the cross-session average number of reward cells for each mouse. **c,** Tuning curves of 4 representative reward cells are shown. First column: Place field. Second column: Vectorized place field. Third column: Trial-by-trial calcium activity (top panel: when mouse approaches reward from left - after touching the left screen; bottom panel: when mouse approaches reward from right - after touching the right screen). The first vertical white line shows the onset of the reward, and the second white line shows the end of the reward consumption. Trials are sorted by their reward consumption duration. Fourth column: average calcium traces with shaded SEM across trials—Blue for those with left screen choice and black for trials with right screen choice. **d,** Average neuronal activity (average deconvolved traces) during ITI vs task are compared. The top panel is for one mouse (mouse #6) across sessions (each point being a reward cell), and the bottom panel is across mice (each point representing one session - averaged across all reward cells for that given session). Reward cells are more active during the task than ITI (See Extended Data Fig. 9). **e,** The percentage of identified reward cells for each session vs time (session) is plotted. The negative correlation of the percentage of the reward cells and time reveals that the recruitment of reward cells decreases over time. The correlation coefficient = −0.52 (P-value = 0.0096). **f,** The percentage of reward cells for each session vs mouse performance is plotted. % reward cells only weakly correlate with mouse performance. The correlation coefficient = −0.15 (P-value = 0.48). **g,** Correlation analysis across all mice (n = 8) indicates that the percentage of recruited reward cells in the hippocampus is negatively correlated with time, decreasing over time during learning and remaining unaffected by performance. Wilcoxon signed-rank test P-value = 0.0078. **h,** A linear model shows that time rather than performance explains most of the variance in the dynamics of % reward cells. The variance explained by time = 0.34 ± 0.07; the variance explained by performance = 0.05 ± 0.02.

### Pre-reward encoding increases across learning

In the previous section, we demonstrated that hippocampal reward encoding diminishes over time at both the population and single-cell levels. In this section, we apply the same methodology used to measure hippocampal reward representation to quantify the evolution of hippocampal encoding of pre-reward moments. Specifically, our study identifies and examines two critical task-related events that occur prior to reward delivery: 1) Screen: during the choice phase two white cues are shown, and the mouse touches the mismatched to sample screen to make its choice. Here we focus on the representation of a window of [-150, 150] ms around touching the screen as screen representation. and 2) Reward approach: the period between a correct choice and the onset of the reward consumption, when the mouse runs from the screen to the reward port. Using the same methods used for the reward analysis, we conducted both population-level and single-cell-level analyses to measure the encoding of screen and reward approach epochs separately to observe how the hippocampal representation of these features evolves over time.

For screen encoding during choice, we observe distinct subpopulations of neurons encoding left and right screen touches (Fig. 4a-b). These subpopulations consistently respond whenever the mouse interacts with each screen. Similar to our analysis for reward, we used a 5-fold decoding analysis to predict screen events from latent neuronal representations. For each fold, the Mutual Information (MI) between predicted and actual screen events was calculated, and the average MI across the 5-folds was considered as the information content of screen representation. Correlating the information content of screen representation with time (session) and mouse performance reveals a positive correlation for both factors (Fig. 4c). A linear model assessing the screen information content in relation to time (session) and mouse performance indicates that both factors significantly contribute to explaining the variance in the dynamics of the screen representation (Fig. 4d). A similar analysis for reward approach encoding suggests the same trend for the information content of reward approach representation (Fig. 4e-h).

**Fig. 4.**
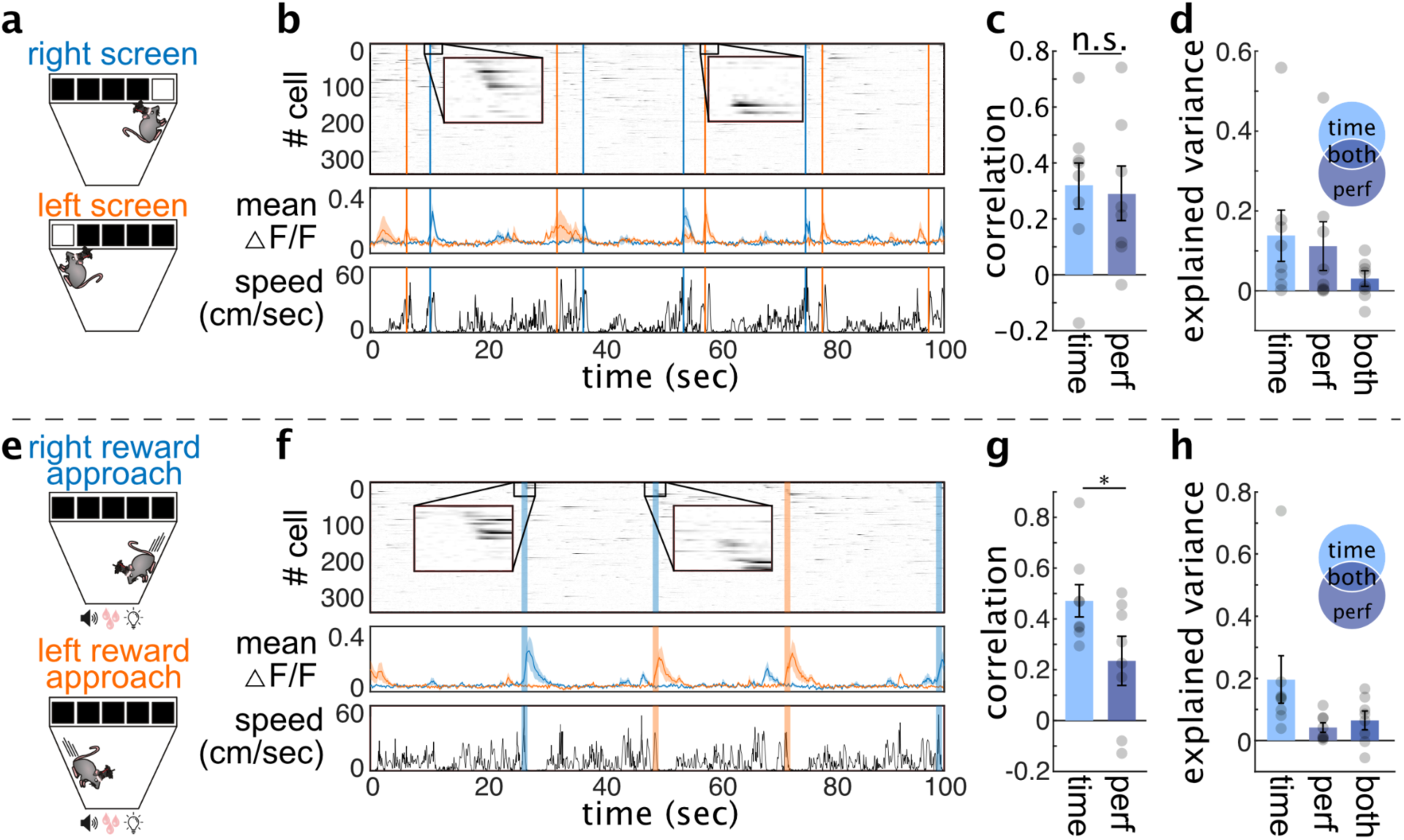
Increased representation of screen and reward approach coding during learning. **a,** The schematic of screen moments; the top panel shows the poke to the right screen, and the bottom panel shows the poke to the left screen. **b,** Top panel: Raw calcium traces of 350 recorded cells are shown. The blue and red vertical lines indicate poke moments to the right and left screens, respectively. The first 26 cells are identified as screen cells and are sorted based on their peak activity around the poke moment. Left and right screens recruit distinct subpopulations of neurons: cells 1 to 15 are active at the right screen, and cells 16 to 26 are active at the left screen. Middle panel: The average calcium activity for each left and right screen cell is plotted. Bottom panel: The speed profile of the mouse is displayed. **c,** Using a similar approach to the quantification of the information content of reward representation presented in Fig. 2, here, we calculate the information content of the screen representation to see how it evolves with time and performance. Correlation analysis shows that screen representation positively correlates with time (session) and performance. Wilcoxon signed-rank test for P-value = 0.3125. **d,** A linear model of screen MI as a function of time (session) and performance demonstrates that both time and performance play significant roles in explaining the variance of dynamics of screen MI. The variance explained by time = 0.14 ± 0.06; the variance explained by performance = 0.12 ± 0.05; the variance explained y covariance of time and performance = 0.03 ± 0.02. The second row (panels **e-h**) repeats the same analysis for a reward approach moment - the time from the screen to the reward. The same analysis shows that the reward approach representation increases with time and performance, aligned with screen representation and in opposition to reward representation. Wilcoxon signed-rank test for panel **g** P-value = 0.0391. **h,** The variance explained by time = 0.19 ± 0.08; the variance explained by performance = 0.04 ± 0.02; the variance explained y covariance of time and performance = 0.07 ± 0.03.

Additionally, we examined the evolution of screen and reward approach representation at a single-cell level. To do this, we used a shuffle-control procedure to identify each of the cell types (Fig. 5a and Fig. 5f). On average, we observed that 7.5 ± 0.7% of the cells were identified as screen cells (Fig. 5a, 5b) and 5.7 ± 0.7% of the cells were identified as reward approach cells. Single-cell plots show screen and reward approach cells are behaviorally modulated (Fig. 5c-d, 5h), meaning that their activity depends on wheter they are on the right or left side of the chamber. The percentage of identified cells for both screen and reward approach cells show a positive correlation with both time and mouse performance (Fig. 5c and Fig. 5h). A linear model reveals that both time and performance significantly contribute to the dynamics of recruitment of screen and reward approach cell types (Fig. 5e, 5j).

**Fig. 5.**
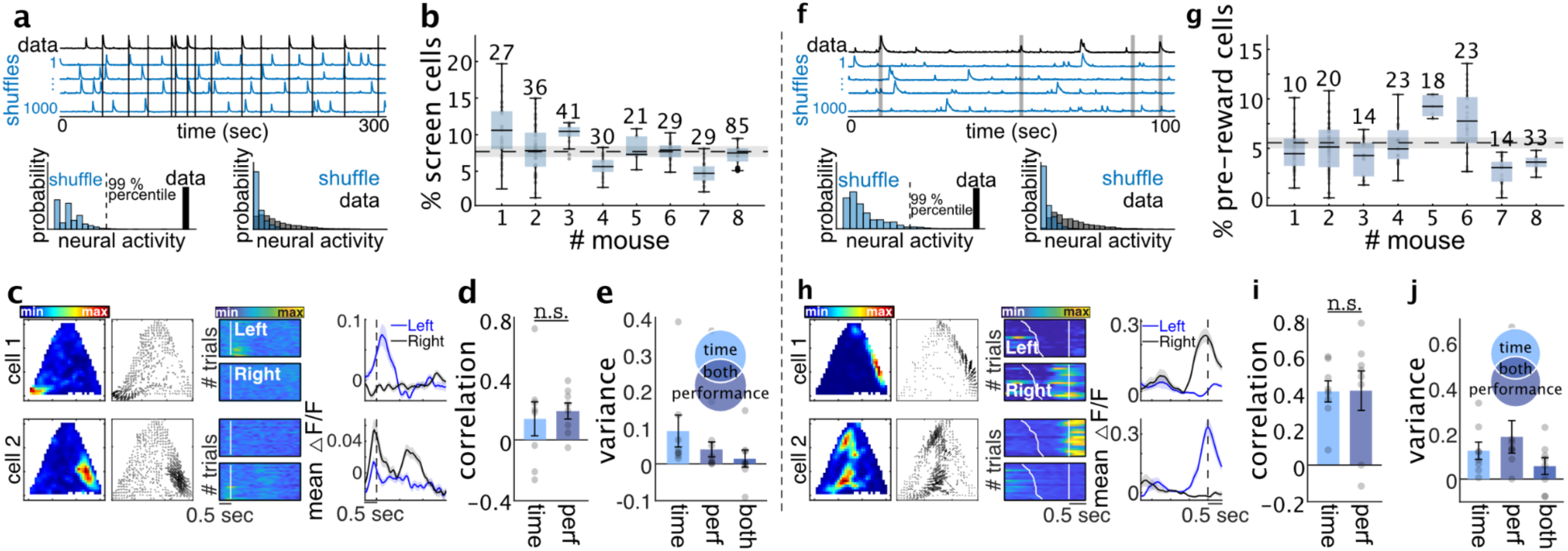
Increase of screen and reward approach cell recruitment during learning. **a,** The same shuffle control procedure as described in Figure 3a is used here to identify screen cells. Neuronal activity 150 ms before and after the screen poke is used to assess neuronal activity at the screen. The panel at the bottom right compares shuffle and control neural activity across all screen cells recorded across sessions and mice. **b,** Percentage of identified screen cells across mice. Each point represents one session, and the numbers above each box show the cross-session average number of reward cells for each mouse. **c,** Tuning curves of 2 representative screen cells are shown. First column: Place field. Second column: Vectorized place field. Third column: Trial-by-trial calcium activity for the left (top panel) and right (bottom panel) screen. The white indicates the time the mouse pokes the screen. Fourth column: average calcium traces with shaded SEM within each group of the left (blue curve) and right (grey curve) screen pokes. The vertical dashed line is the screen poke time. **d,** The correlation between the percentage of screen cells and time and performance across mice is depicted. Both time and performance positively correlate with the percentage of screen cells. Wilcoxon signed-rank test P-value = 0.1953. **e,** Linear modeling of screen cell percentages shows that both time and performance significantly contribute to explaining the variance in the evolution of the percentage of screen cells. The variance explained by time = 0.09 ± 0.04; the variance explained by performance = 0.04 ± 0.02. Panels **f-i** show the same analysis as for panels **a-e** but for reward approach cells. Cells are identified as reward approach cells if their activity from screen to reward is more than chance. Panel **h** column three: trial-by-trial calcium traces. The first white line shows the screen poke and the second vertical white line indicates the onset of the reward. Similar to screen cells, reward approach cells also show an increase with time and performance. Wilcoxon signed-rank test P-value for panel (i) = 0.1953. **j,** The variance explained by time = 0.13 ± 0.04; the variance explained by performance = 0.20 ± 0.08.

Finally, we examined the amplitude of the calcium response of reward approach cells when the mouse approaches reward vs. incentive (a smaller portion of reward given to mice during the delay to keep the mouse engaged in the task) and observed that reward magnitude significantly modulated reward approach cells activity (Extended Data Fig. 10).

### Backward shift of reward coding during learning

In the previous sections, we showed a gradual decline in reward encoding alongside a corresponding increase in the encoding of reward-predicting cues over weeks of extended experience. One potential mechanism to account for these changes is that reward-encoding cells may progressively shift their activity backward to encode the time periods that predict the reward. To investigate this hypothesis, we tracked neurons identified as reward cells in at least one of the first five recording sessions. We included reward cells that we could track for at least five following sessions, resulting in 194 reward cells across all mice. We analyzed the timing of the activity of these neurons relative to reward onset across sessions (Fig. 6a). Our data reveal that a significant number of reward cells exhibit a backward shift from reward toward reward approach and screen cues – termed backward shifting reward cells (Fig. 6a-b, Extended Data Fig. 11). Looking at the difference between peak timing and session number across all backward shifting reward cells shows a clear negative correlation (Fig. 6c). A shuffle-control method was employed to identify the backward shifting reward cells, showing that approximately 23% (45 out of 194 cells) of reward cells exhibit this phenomenon, compared to a chance level of 5% (Fig. 6d). A substantial portion (56%, 25 out of 45 cells) of backward shifting reward cells shifted enough to be classified as screen or reward approach cells in later sessions (Fig. 6b and Extended Data Fig. 11). The temporal progression of these cells was quantified by correlating peak activity timing relative to reward onset with session number (Extended Data Fig. 11). As a control, we performed the same analysis for screen cells and found only 6% of the screen cells are identified as backward shifting (chance level 5%) (Fig. 6d). The correlation between peak activity timing and session number for all reward cells demonstrates a negative trend, which is more pronounced in backward shifting reward cells (Fig. 6e). To determine the backward shifting rate, we measured the slope of peak timing versus session number, finding an average time scale of approximately ∼0.1 seconds per session – indicating it takes around 10 sessions for a reward cell to exhibit a 1-second backward shift (Fig. 6f). These time scales are comparable to the temporal distance between cue and reward and the number of sessions typically required for a mouse to learn the task.

**Fig. 6.**
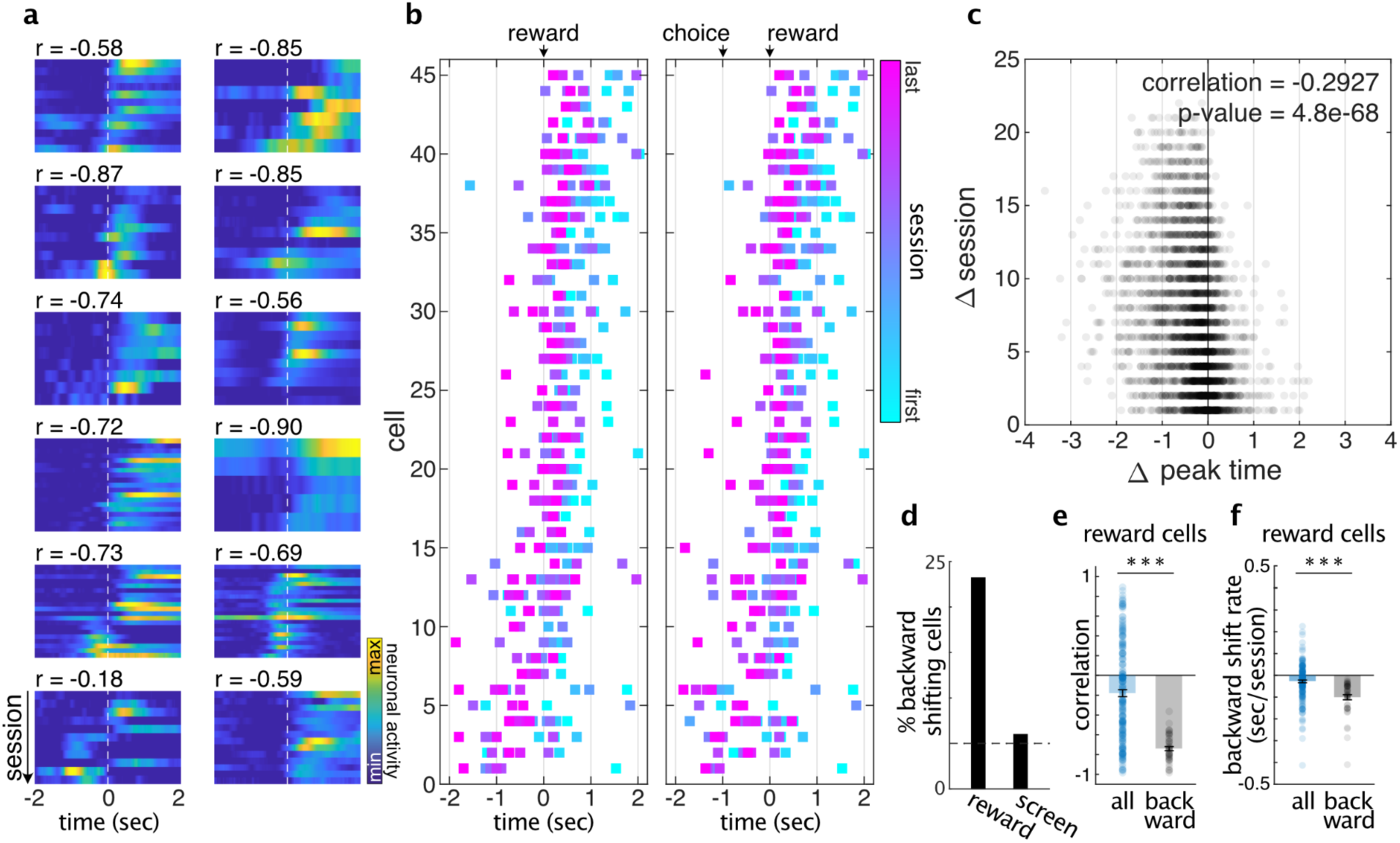
Weeks-long backward shift of reward coding during learning. **a,** Twelve representative backward-shifting reward cells are plotted. The number above each panel shows the correlation between the timing of peak activity in each session and the session number. The negative correlation demonstrates backward shifting. **b,** Left panel: The timing of peak activity relative to the reward onset across sessions for all backward shifting reward cells are presented. The gradient of colors represents the backward shift of peak activity. In the right panel, we normalized the time between choice and reward to one second, so time −1 represents the choice. Cells are sorted based on their mean peak timings. **c,** The difference between peak time and session number for each session is shown. **d,** The percentage of reward cells that passed the backward shifting criteria is ∼23% (45 out of 194 reward cells tracked across more than five sessions). As a control, we repeated the same analysis for screen cells and found out only ∼6% of screen cells passed the backward shifting criteria (8 out of 129 screen cells tracked for more than five sessions). The dashed line shows the chance level is at 5%. **e,** The correlations between peak time and session number for all reward cells (blue, n = 194) and for backward-shifting reward cells (black, n = 45) are depicted. Kolmogorov-Smirnov test P-value = 2.0351e-15. **f,** The backward shifting rate (slope of session number vs. peak timing) for all reward cells (blue, n = 194) and backward shifting reward cells (black, n = 45) are plotted. We observe a backward-shifting slope of ∼0.1 sec/session - meaning a backward shift of ∼1 sec over ten sessions. Kolmogorov-Smirnov test P-value = 6.5824e-10.

## Discussion

We combined large population recordings of mouse CA1^37^ neurons with an automated touchscreen reward-based task^43,44^ to investigate the long-term dynamics of reward encoding in the hippocampus. We observed significant gradual changes in the representation of the task’s salient features across extended experience—mainly during the reward delivery and in responses to cues that predicted reward. Our data revealed a reduction in reward signal and an increase in the response to the cues that anticipate the reward delivery. This was further supported by detecting cells that are initially tuned to the reward and gradually shifted backward to encode aspects of the task that are reward predictors.

Our data supports the notion that the hippocampus does more than representing the animal’s location; it is learning to predict the animal’s overall state, with reward being a crucial component of that state. This aligns with recent models that view place cells as encoding predictions of future states enabling the hippocampus to efficiently anticipate state-state transitions^9^. Building on previous studies of how reward learning influences hippocampal neuronal representation^13–19^, we demonstrate that this modulation is highly structured and results in a reward-predictive map.

Previous studies have revealed that hippocampal place fields move towards goal locations early in learning, likely contributing to what others have observed as an over-representation of rewarded locations^15,16^. Here, we also observe an over-representation of the reward location (Extended Data Fig. 6) after 2-3 weeks of pre-training, in which animals are trained to touch the touchscreen for reward. Other work has shown a backward shift of hippocampal place fields, independent of reward locations, on a faster, within-session timescale^58^. Together, these outcomes suggest that the hippocampal representation initially over-represents rewarded locations and is followed by a slower, weeks-long shift to represent the cues that predict these rewards.

Another interpretation of our results could be that the reward signal in the hippocampus may act as a teaching signal, encoding reward prediction error, the difference between expected and actual rewards. The dopaminergic system in the brain is known for reward computation^59–61^, specifically providing a neuronal basis for learning through reward prediction error^1,62^, with Temporal Difference Reinforcement Learning (TDRL) having profoundly shaped our understanding of dopaminergic reward coding^61,63–66^, a concept that has also influenced our understanding of hippocampal physiology^67^. Prevalent implementations of TDRL make two key predictions about reward prediction error coding, which have been observed in dopaminergic systems and in the hippocampus in the current study: 1) a gradual decrease in reward response coupled with a gradual increase in response to reward-predicting cues during learning^1,62^, and 2) a gradual backward temporal shift of the error signal from reward to cues during learning^68^.

While our findings align with certain predictions of TDRL, additional controls would be necessary to conclusively interpret the hippocampal reward signal as reflecting reward prediction error. For example, TDRL theory predicts a suppression of the reward prediction error signal when an expected reward is omitted, a phenomenon observed in VTA dopamine neurons^1,62^. However, our current task design does not allow us to explore this aspect, as mice in our study do not check the reward port during incorrect trials due to the task structure. This limitation precludes a thorough examination of the reward cells’ response in such scenarios.

In conclusion, our study uncovers a dynamic and organized backpropagation of hippocampal reward representations during the learning process. Far from serving as a mere spatial map, the hippocampus exhibits predictive coding, progressively tuning its representation to anticipate future rewards. These insights advance our understanding of the hippocampal role in learning, highlighting its crucial contribution to the brain’s overarching objective of forecasting and optimizing future rewards.

## Acknowledgments

We thank Daniel Aharoni for his guidance in using the UCLA miniscope. We thank the members of the Brandon laboratory for discussions and providing inputs during the analysis of the data, in particular, Zaki Ajabi and J. Quinn Lee. We thank Mihaela Iordanova, Blake Richards, Etienne JP Maes, J. Quinn Lee, Harshith Nagaraj, Zeeshan Haqqee, and Apoorv Sharma for insightful comments on the first draft of this manuscript. We thank Adrien Peyrache and Blake Richards for their valuable advice and guidance on data analysis. This work was supported by funding from Fonds de Recherche du Québec – Santé (FRQS) postdoctoral fellowships awarded to A.N. and C-A.M., CIHR Project Grants #463403 and #480510 to M.P.B, and a Core Facilities and Technology Development grant from Canada First Research Excellence Fund, ‘Health Brains for Health Lives’ to S.W. and M.P.B.

## Data and Code Sharing

All data and source codes used in the current study will be made available upon publication.

## Material and methods

### Subjects

Eight naïve male mice (C57BL/6 mice, Charles River) were housed individually and maintained a 12-hour light/dark cycle at 22°C and 40% humidity with water ad libitum. All experiments were carried out during the light portion of the light/dark cycle and were in accordance with McGill University and Douglas Hospital Research Center Animal Use and Care Committee (protocol # 20157725) and in accordance with Canadian Institutes of Health Research guidelines.

### Surgeries

Animals underwent three surgeries under isoflurane (1.5-2%, vol/vol). In addition, carprofen (10ml/kg) and saline (0.5ml) were administered at the beginning of each surgery. We injected 400nl of either AAV9.syn.GCaMP6f.WPRE.SV40 (University of Pennsylvania Vector Core, 3.26e14 GC-ml) diluted 1:1 with PBS or AAV5-CaMKII-GCaMP6f.WPRE.SV40 (Addgene, 2.3e13 GC-ml) diluted with PBS 1:2 into dorsal CA1 (−1.8 mm from Bregma, 1.5mm mediolateral, 1.45mm dorsoventral). Two weeks after viral injections, a gradient refractive index (GRIN) lens (Edmund Optics, 1.8mm in diameter, 0.25 pitch, 4.31mm in length) was implanted above the previous injection site. Briefly, a 1.8mm craniotomy above the injection site was done followed by aspiration of cortical tissue directly below the craniotomy. The GRIN lens was lowered to the area of interest and two stainless steel screws were threaded into the contralateral skull. Both the GRIN lens and the screws were fixed with dental cement (C&B Metabond). Silicone adhesive was used to cover the lens until the next surgery. 2-3 weeks after the GRIN lens implant an aluminum baseplate was attached with dental cement to the mouse’s skull and covered with a plastic cap to protect the lens.

### Apparatus

Mice were trained in the Bussey-Saksida automated touchscreen operant chamber (Lafayette Instrument) described here^44,69^. Briefly, this trapezoidal-shaped apparatus features a touch-sensitive LCD computer monitor (12.1-inch screen, 800 × 600 resolution) at one end and a reward collection magazine (20 cm H × 18 cm L × 6-24 cm W) at the other, tapering from the touchscreen to the magazine. The arena’s walls are made of black Perspex, and the floor is perforated stainless steel with a stainless-steel waste tray underneath. The entire setup is housed in a sound- and light-attenuating box equipped with a house light, a tone generator, and a ventilating fan.

Above the arena, a house light (3 W) and a video camera are mounted. A peristaltic pump is positioned centrally behind the touchscreen unit to deliver the liquid reward; in our experiment, we used strawberry-flavored milkshake (Québon, Agropur, CA) as the food reward. An infrared (IR) beam detects entries into the reward-delivery magazine, which is equipped with a light and a small speaker. Additionally, two IR beams cross the arena to detect locomotor activity.

To minimize unintended screen touches and to demarcate screen response locations, a black Perspex mask with five response windows (each comprised of a 4 × 4 cm square aperture, 1.5 cm above the grid floor) covered the touchscreen to reduce incidental touches. The task schedules were designed, managed, and events recorded using Whisker Server and ABETTII software (Campden Instruments).

### Behavior

Animals were food-deprived until they reached 85-90% of their original weight. Before starting the Trial-Unique Nonmatching-to-Location task, the mice underwent several behavioral training stages as previously described^45^.

### Pretraining

Initially, the mice were habituated to human handling in the touchscreen chamber room for 3 days. Following this, they were acclimated to the chamber itself, with rewards presented in the reward tray. They could progress to the next stage once they finished the reward within 20 minutes, typically within 1-2 days.

After the habituation phase, the mice were trained to touch the screen when a white square stimulus was presented pseudo-randomly in one of five possible locations on the screen. A reward was given when the mouse touched the screen while the sample was displayed. The mice progressed to the next stage after completing 30 trials within 60 minutes.

The next stage required the mice to touch the white square on the screen to receive a reward, with the same completion criterion of 30 trials within 60 minutes. Subsequently, the mice had to learn to initiate trials by moving to the back of the chamber and breaking the IR beam near the reward magazine.

In the final pretraining stage, a touch to blank windows resulted in a 5-second timeout, signaled by the illumination of the house light. Correction trials, which repeated the same trial after a 5-second inter-trial interval (ITI), were administered until the mouse made a correct response. However, these correction trials were not included in the performance calculation. Reward collection initiated a 15-second ITI before the next trial began.

### Task

The TUNL task consists of two phases: the sample phase, an encoding phase, where the mouse learns the location of the cue, and a retrieval phase, where it has to remember the cue location and choose the non-matching one. The first stage of TUNL training is designed to teach the non-matching rule, requiring the mouse to identify the novel location as the correct choice. (Fig. 1F), During the sample phase, one of five locations on the touchscreen is illuminated. Upon a nose poke to this location, the mouse is directed to the back of the chamber by the illumination of the reward tray (an 800 millisecond (ms) pulse delivering 20 μL of milkshake) and an auditory tone.

The delay length is initially maintained at 2s during learning of the non-matching rule and is then increased by an increment of 2s during specific probe trials. Once the back IR beams are broken following the delay period, the original sample and a novel correct location are presented simultaneously on the touchscreen.

If the mouse makes an incorrect response to the original sample location, a correction trial loop is initiated until the correct response is made. Correction trials are repeated presentations of the same sample and choice locations following an incorrect response. Correction trials are not included in the percentage of correct responses. Mice were trained until they reached an average of 70% correct over two sessions of 36 trials. Once the mice reached the criterion for trials with a 2-second delay, they progressed to trials with a 4-second delay, and so forth.

### Data acquisition

*In vivo* calcium videos were recorded using a UCLA miniscope^37^ (v3; miniscope.org) equipped with a monochrome CMOS imaging sensor (MT9V032C12STM, ON Semiconductor). This sensor was connected to a custom data acquisition (DAQ) box (miniscope.org) via a lightweight, flexible coaxial cable. The DAQ box was linked to a PC using a USB 3.0 SuperSpeed cable and operated with Miniscope custom acquisition software (miniscope.org). Behavioral videos were recorded with an infrared camera positioned above the touchscreen to monitor mouse behavior. The DAQ simultaneously acquired behavioral and cellular imaging streams at 30 Hz as uncompressed avi files and all recorded frames were timestamped for post hoc alignment. Touchscreen chamber also provides task-related information such as trial initiation timing, nose pokes to the screen, and reward onset, and other task related information. A touchscreen chamber timestamp is also provided for follow-up alignment with neuronal and behavioral data.

## Data analysis

### Pre-processing of calcium and behavioural data

We have used a UCLA miniscope to simultaneously record several hundreds of neurons in a freely-moving mouse^37^. This provides the possibility of monitoring hundreds of neurons that are located inside of our field of view. The output of this recording in our experimental setup is a video with 30 frames/sec temporal and 2-3 *μm* spatial resolution. The temporal resolution is sufficient to capture the slow dynamics of calcium transients and the spatial resolution is sufficient to capture the cell bodies. The main steps for analyzing the calcium recording videos are as follows: 1) Within-session motion correction to address small displacements and shakes during recording; 2) Detecting cell bodies; 3) Extracting calcium traces for each cell body by measuring the average fluorescent emission from the detected cell body; 4) Inferring the likelihood of spikes from the raw calcium traces.

Calcium imaging data were preprocessed prior to analyses via a pipeline of open-source MATLAB (MathWorks; version R2021a) functions to correct for motion artifacts^39^, segment cells and extract transients^39,41,42^. A first-order autoregressive model is used to infer the likelihood of spiking events through the deconvolution of the transient trace through ^42^. The resulting time series are used to measure the ‘firing rate’.

DeepLabCut, a deep-learning based tool for pose estimation, is used to track multi-part of the mouse^70^. The tracking is used to estimate position, heading direction, speed, and other behavioral features.

### Identification of cell types

To identify each cell type (reward, reward approach, and screen cells), the averaged neuronal response of each cell to each of the three features was calculated. Averaged neuronal activity at reward: average deconvolved traces during the reward consumption period. Averaged neuronal activity at reward approach: average deconvolved traces between the correct choice and onset of the reward. Neuronal activity at the screen: average deconvolved traces at a window of [-150 ms, 150 ms] around screen pokes during choice period. The averaged neuronal activity for each cell is compared to the distribution of averaged neuronal responses made by 1000 circular shuffles. Cell types were identified as those whose neuronal activity exceeded the 99th percentile of the corresponding shuffled distribution.

### Naive Bayes spatial decoding

To decode animals’ positions from calcium traces within each session, we divided the spatially binned position (using spatial bins of 1 cm along each of the axes) and our deconvolved calcium traces into 5-fold splits. The binned positions were converted into a one-hot vector. Using the Gaussian Naïve Bayes method from the scikit-learn Python library, we predicted positions on the withheld data based on maximum likelihood estimation, as follows:

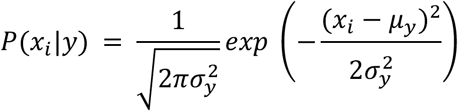

where *x_i_* corresponds to the predicted position at *i* and *y* to the respective calcium traces. We assumed a flat prior (equal likelihood at all positions) and the scikit-learn default σ value of 1e-9. The decoded position is assigned to the bin with the highest probability. The decoding error was then estimated as the Euclidean distance between the animal’s predicted and actual spatial bin position on withheld data.

### Mutual Information (MI) content of task features

CEBRA embedding is used to project the deconvolved neuronal traces into a 32-dimensional latent space^57^. We set all parameters as default and used the same set of parameters across all sessions. The projection was done in a self-supervised fashion when no behavioral or task-related information was involved. The feature of interest (screen, reward approach, or reward) was presented as a binary vector when for each frame if the mouse is in that condition it is 1 otherwise it is 0. A 5-fold cross-validation is implemented where for each fold one fold of the data is held out, a scikit-learn-based linear decoder is trained on the latent presentation to decode the class of the binary target. The Mutual Information (MI) (using sciki-learn) between the predicted and actual target for the held-out data is calculated. The averaged MI across five folds is considered the MI for each recording session.

### Linear modeling of Information content

To compute the contribution of each feature (time and mouse performance) in the evolution of hippocampal representation, we have a Matlab function, fitlm, to model each of the measures of interest denoted by Y, by time and performance. The contribution of each of the features is measured by their contribution to the explained variance of Y:

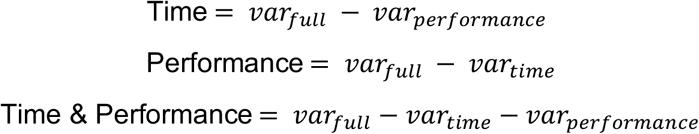

*var_time_*: Model’s explained variance when Y is modeled only by time.

*var_performance_*: Model’s explained variance when Y is modeled only by performance.

*var_full_*: Model’s explained variance when Y is modeled by both time and performance.

For a visual demonstration look at the Venn diagram in Fig. 2i.

### Identification of backward-shifting reward cells

Averaged calcium traces relative to the onset of the reward is calculated for each of the reward cells. The correlation between the session number and the time of peak activity is calculated. A shuffle distribution is calculated by correlating the time of peak activities and the shuffled session numbers. A cell is identified as a backward-shifting reward cell if its correlation is less than the 5th percentile of shuffle distribution. This criterion sets the chance level to be 5%.

### Rate maps

To calculate the rate map for each neuron we binned the x and y axis each into 30 bins. The rate value assigned to each bin was simply calculated by the sum of neuronal activity (deconvolved traces) normalized by the time the mouse spent in that bin. For visualization, we used a Gaussian filter of size 5×5 bins and sigma = 1 bins.

### Place cell’s identification

We computed the spatial information of all cells using the unsmoothed-event-rate map of each cell, as previously described^71^.

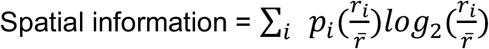

Where *p_i_* is the probability of the mouse being in the *i^th^* bin (time spent in *i^th^* bin/total running time); *r_i_* is the Ca*^2+^* event rate in the *i^th^* bin; and *r* is the overall Ca*^2+^* event rate. We then performed 1000 distinct shuffles of animal locations during Ca2+ events and calculated the spatial information for each shuffle. Cells with spatial information higher than that of 99% percentile of their shuffles were identified as place cells (Extended data Fig. 3).

### Measuring the size of the place fields

After identifying place cells and determining their rate maps, we masked these rate maps by setting all bins with values below the 90th percentile of values across all bins to zero. This operation creates islands of non-zeros surrounded by bins of zero-value. We used the Matlab function `bwconncomp` to detect these islands and used `regionprops` to calculate the area of each of these islands. The island with the maximum area was used to calculate the place field size.

The size of the place field was then calculated as:

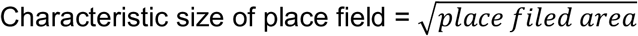

### Classification decoding analysis

A linear support vector machine classifier from MATLAB (fitcecoc) was used to decode contextual information in the task. For example, we want to decode the correctness of the trial at different moments of the task. Population vectors of deconvolved calcium traces were used to train and test the classifier. Given the limited number of samples, we employed a leave-one-out approach by training our classifier on all samples except one and testing it on the excluded sample, repeating this process for each sample point. In each iteration, we ensured that the training dataset contained an equal number of samples from each class by randomly downsampling the class with a larger number of samples. For each decoding, we repeated the process five times and averaged the decoding accuracy across these five iterations. The classifier’s decoding performance was compared to the accuracy obtained from shuffled interaction, where the class labels were randomly shuffled.

### Reward over-representation score

To measure the reward over-representation score within each session, we first generated the spatial rate map for all cells. We identified the location of peak activity for each rate map, detecting the spatial bin with the highest firing rate for all cells. A density plot was then created to represent the density of peaks in each spatial bin. This matrix provides a representation of each spatial bin.

The reward over-representation was calculated as the average representation of the 10% of spatial bins closest to the reward port, normalized by the average representation across all spatial bins. In this context, a score of 1 indicates an even distribution of reward representation compared to the baseline, while a value greater than 1 signifies an over-representation of spatial bins near the reward port.

### Statistical analysis

For visualization, we represented error bars (or shaded areas for line plots) using the standard error of the mean (SEM). To compare two distributions, we employed the Two-sample Kolmogorov-Smirnov test via the ‘kstest2’ command in Matlab. For comparisons of a distribution against zero, we utilized the Wilcoxon signed rank test, implemented using Matlab’s ‘signrank’ command. Significance levels for all tests were set at *p<0.05, **p<0.01, and ***p<0.001.

## Author Contributions

M.Y., A.N., and M.P.B. conceptualized the project. A.N. performed surgeries and recordings. M.Y. organized the raw data and did the pre-processing, analysis, modeling, and data visualization. M.P.B. guided and supervised all the stages of experiments and data analysis. M.Y. and C-A.M. wrote the initial draft. M.Y., C-A.M., M.P.B. contributed to editing and revising the paper.

## Competing Interests

The authors declare no competing interests.

---

**Extended Data Fig. 1.**
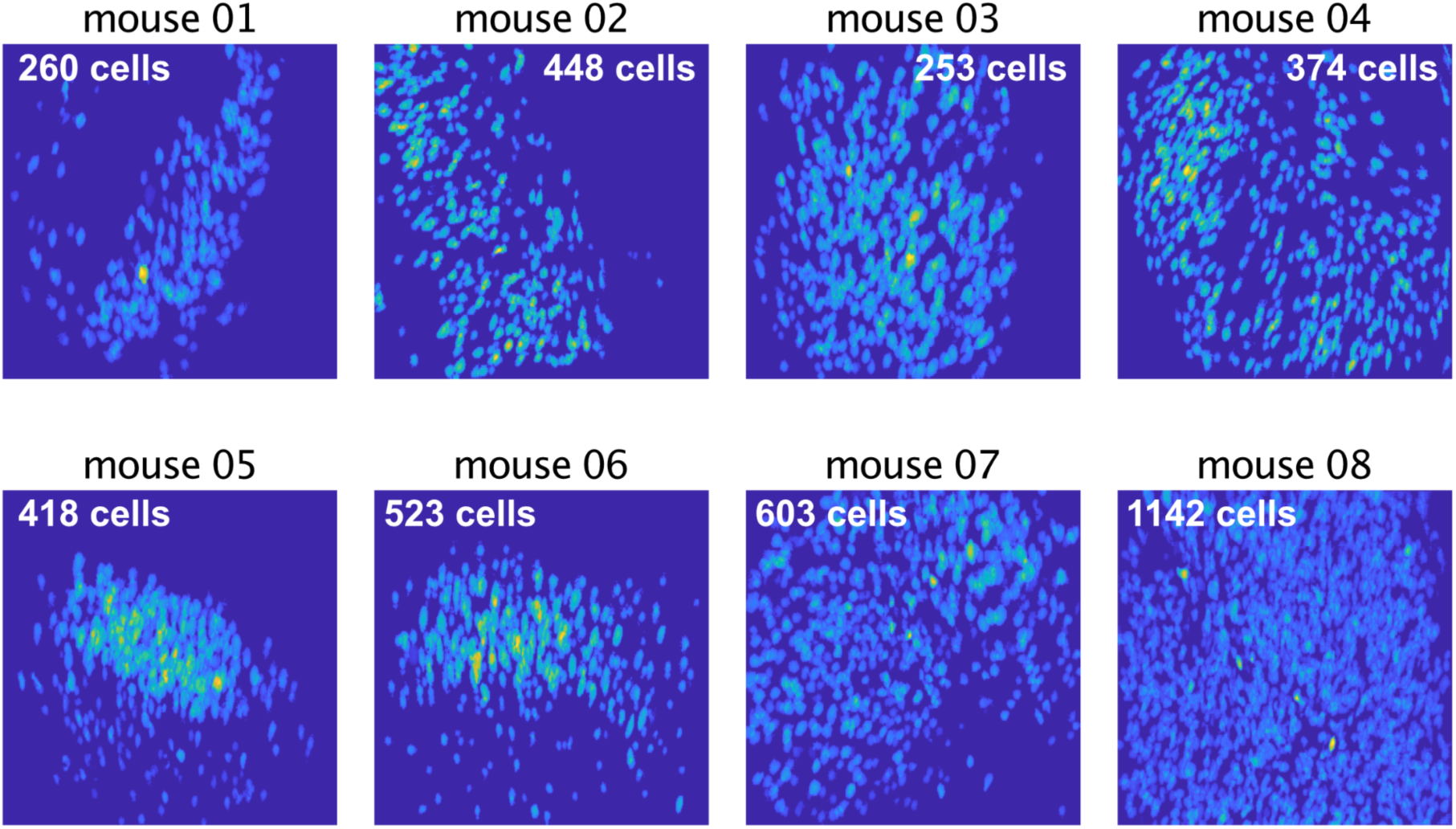
Sample footprints for each of the mice. Footprints of an example session for each mouse are presented. The number in each panel represents the average number of detected cells across sessions for each mouse.

**Extended Data Fig. 2.**
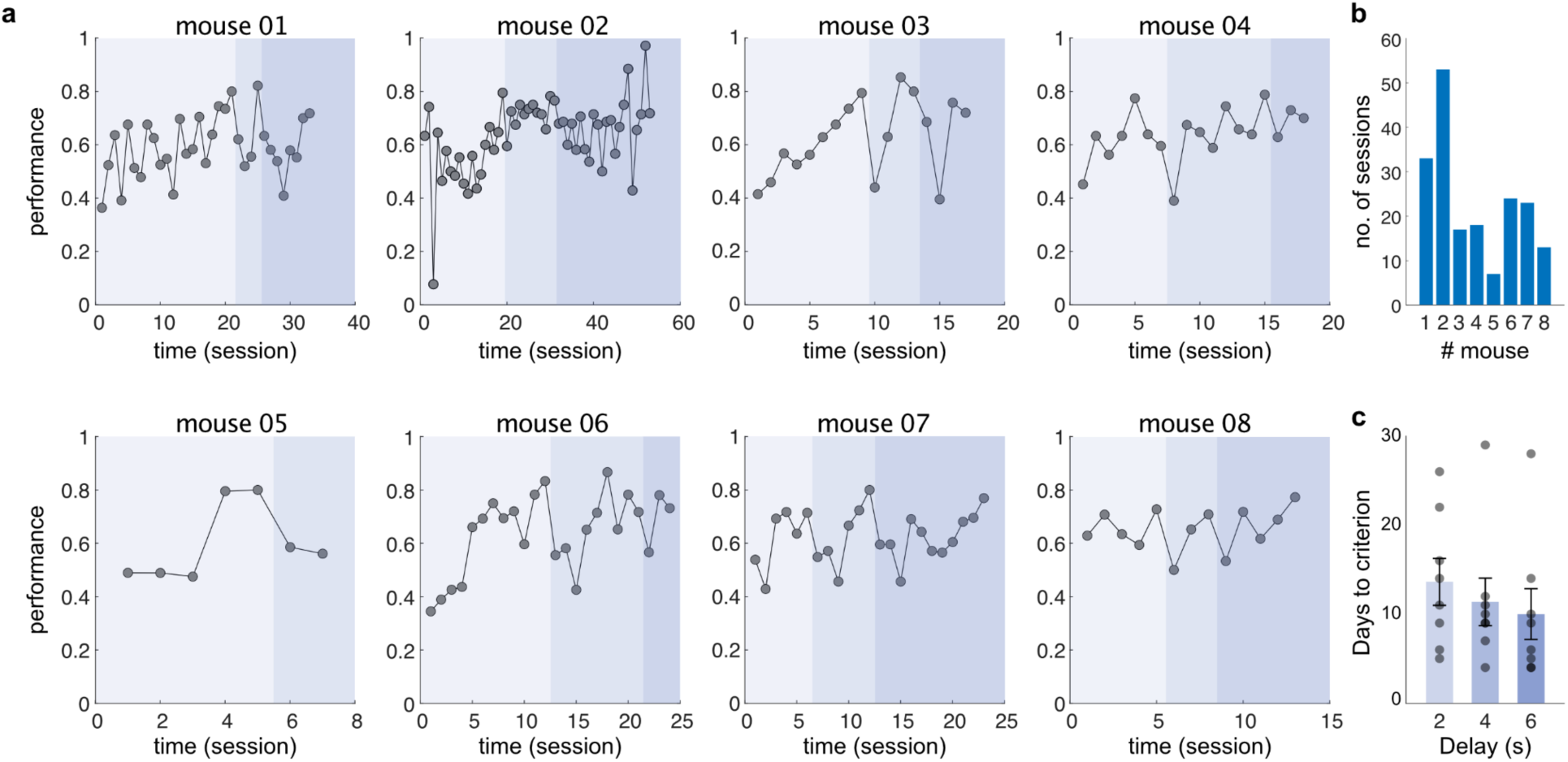
Learning curves. **a,** Learning curves for all eight mice are presented. Performance is defined as the proportion of correct trials. The shades of blue color show different delays from lighter to darker being 2, 4, and 6 seconds. **b,** Number of recorded sessions for each mouse. **c,** The number of days that it takes for each mouse to reach the learning criteria (two consecutive days of performance > 0.7) for different delays. We see longer delays take more time to reach the criteria.

**Extended Data Fig. 3.**
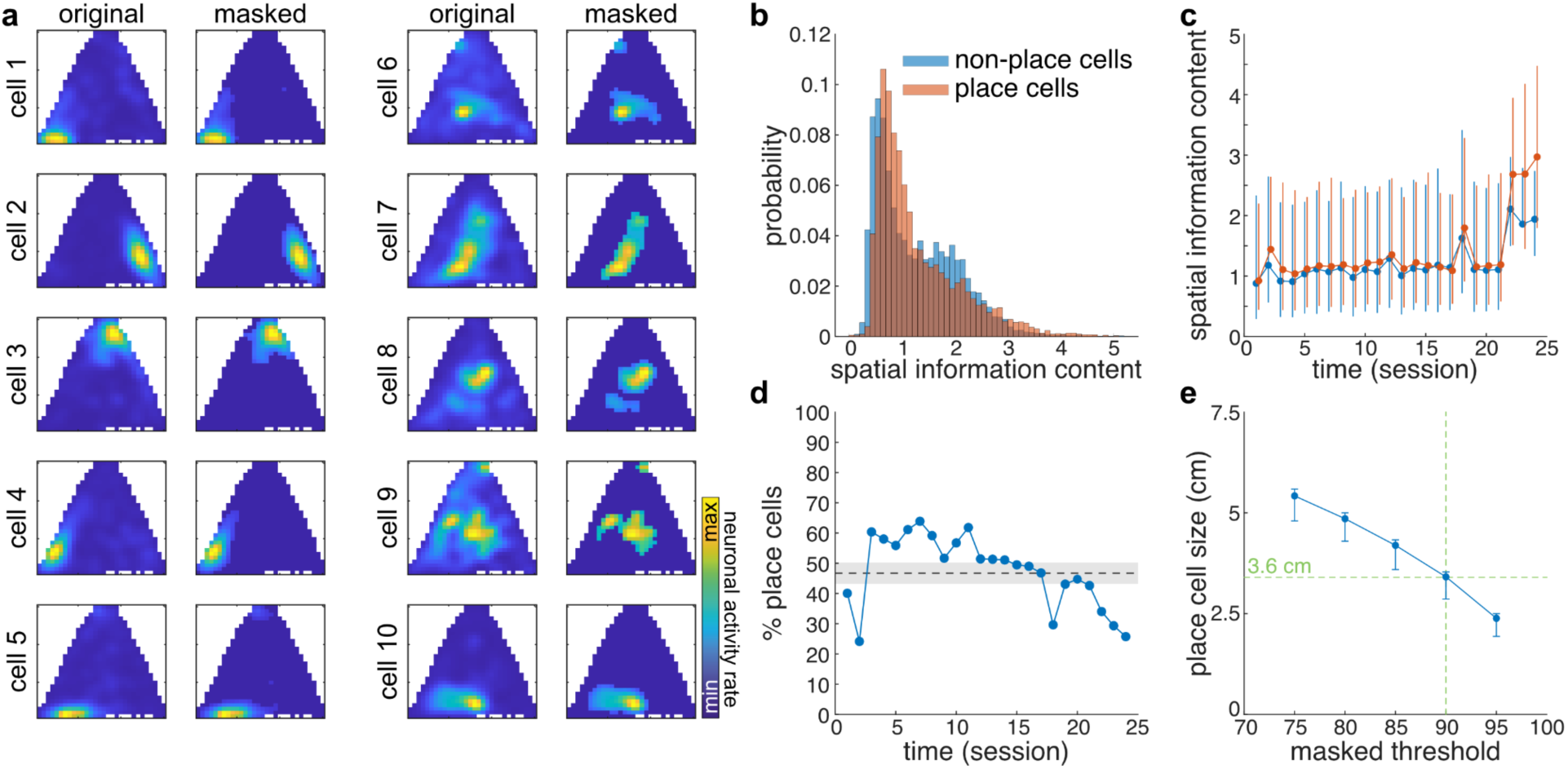
Place field size. **a,** Spatial rate map of ten representative place cells are plotted. For each cell the left panel is the raw rate map and the right panel is the masked rate map (masking has been done by turning all bins with value less than 90% percentile to zero). Place cells are identified if their spatial information content exceeds 99% percentile of distribution of spatial information content for 1,000 shuffled cases. **b,** The distribution of information content for cells that are identified as place cells (orange) vs cells that did not identify as a place cell (blue) are plotted. We have combined all sessions for mouse #6 (as a representative mouse). **c,** The evolution of mean spatial information content across sessions is plotted; color coding is the same as panel b. **d,** Percentage of identified place cells vs time (session) is plotted. In average 47.5 ± 2.5 % of the cells were identified as place cells. **e,** After doing visual inspection we decided to use 90% percentile for masking spatial rate maps to calculate the size of the place field. This choice threshold led to average place cell’s field size = 3.6 cm. However, here, we have calculated averaged place cell’s field sizes for a various amount of threshold from 75% to 95% percentile.

**Extended Data Fig. 4.**
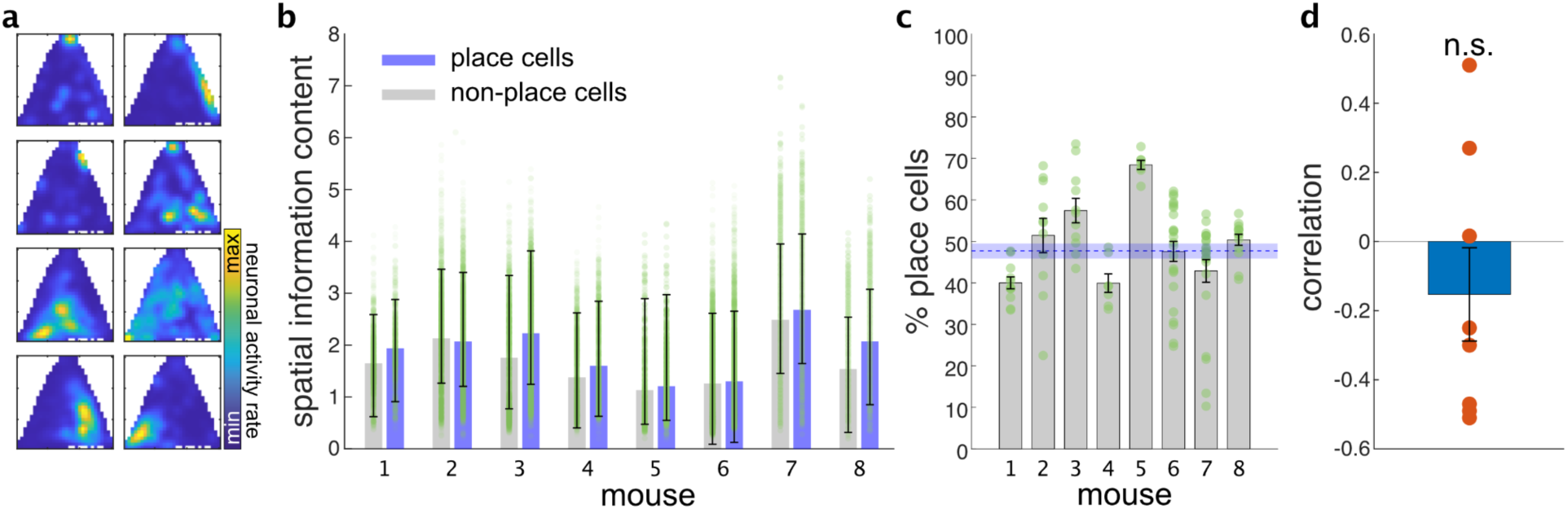
Dynamics of place cell recruitment over time. **a,** Eight place cells are plotted. The place cells are identified using an information theory-based shuffle control procedure as described in the Method section. **b,** Spatial Information content for place cells and non-place cells are compared for all neurons across sessions and mice. Each point represents a cell. **c,** Percentage of identified place cells for all mice is presented. Each point represents a recording session. **d,** The correlation between the percentage of place cells with time (session) is calculated. We do not observe a consistent trend across mice.

**Extended Data Fig. 5.**
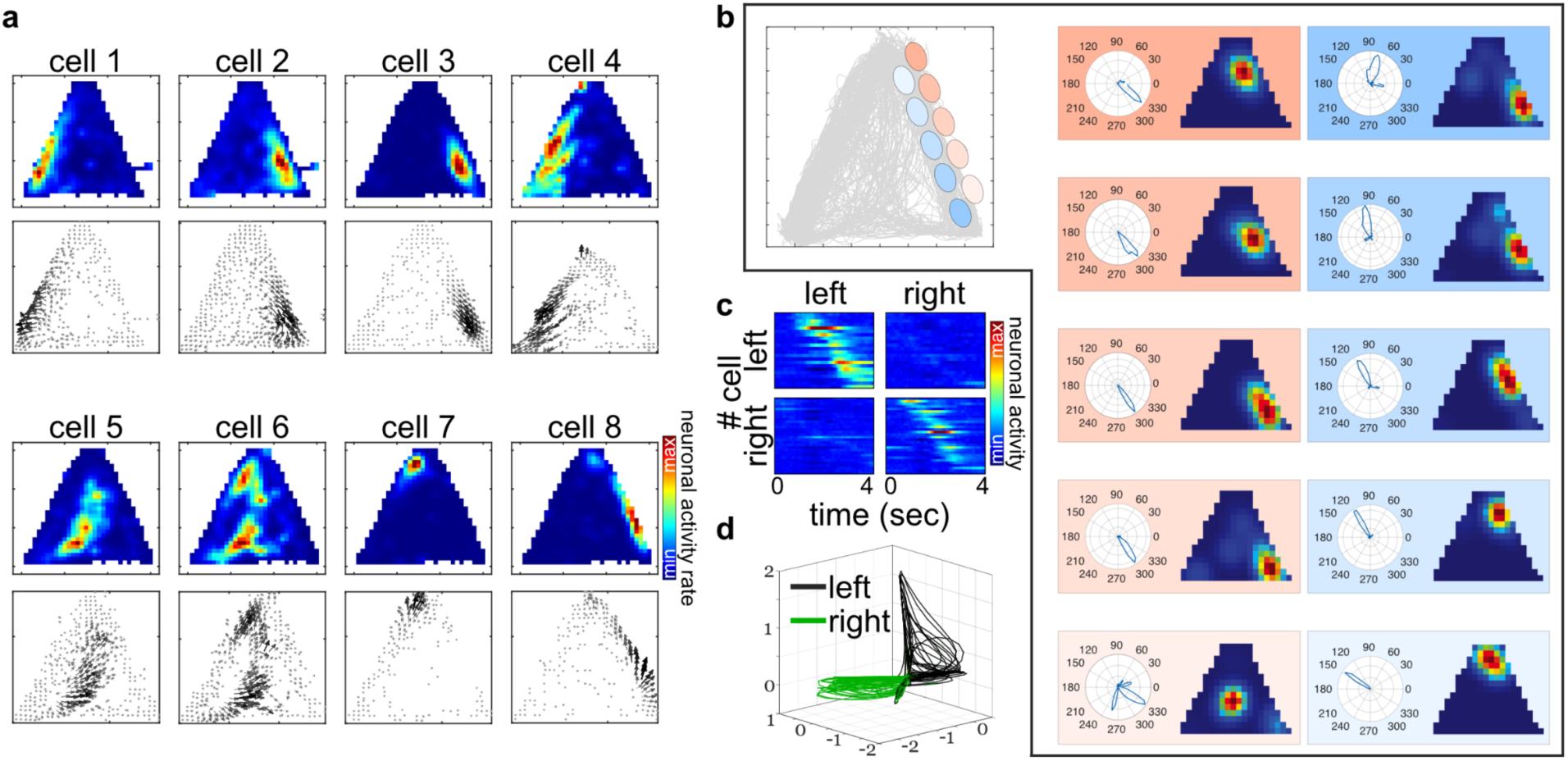
Place cells firing rates are modulated by behavior. **a,** Eight place cells with their rate maps and vectorized rate maps are shown. The vectorized rate maps clearly demonstrate the modulation of neuronal activity by the mouse’s behavior, specifically its heading direction. **b,** Two dominant stereotyped behaviors in this task are the mouse running from the reward area to touch either the right or left screen and then running back to the back of the chamber. A distinct population of neurons encodes different phases of each of these stereotyped behaviors. Here, we visualize 10 place cells encoding different phases of the mouse running from the reward area to the right screen and then running back to the reward zone. Tuning curves include spatial rate maps and heading direction tuning curves. **c,** The sequences of cells for flattened stereotyped behaviors of running to the left and right are plotted, and averaged across trials. Distinct subpopulations of neurons support each of these two behaviors, remaining silent during the other movement. **d,** GPFA dimensionality reduction ^72^ on the entire population for these two patterns of behavior reveals perpendicular neuronal trajectories in the latent state space.

**Extended Data Fig. 6.**
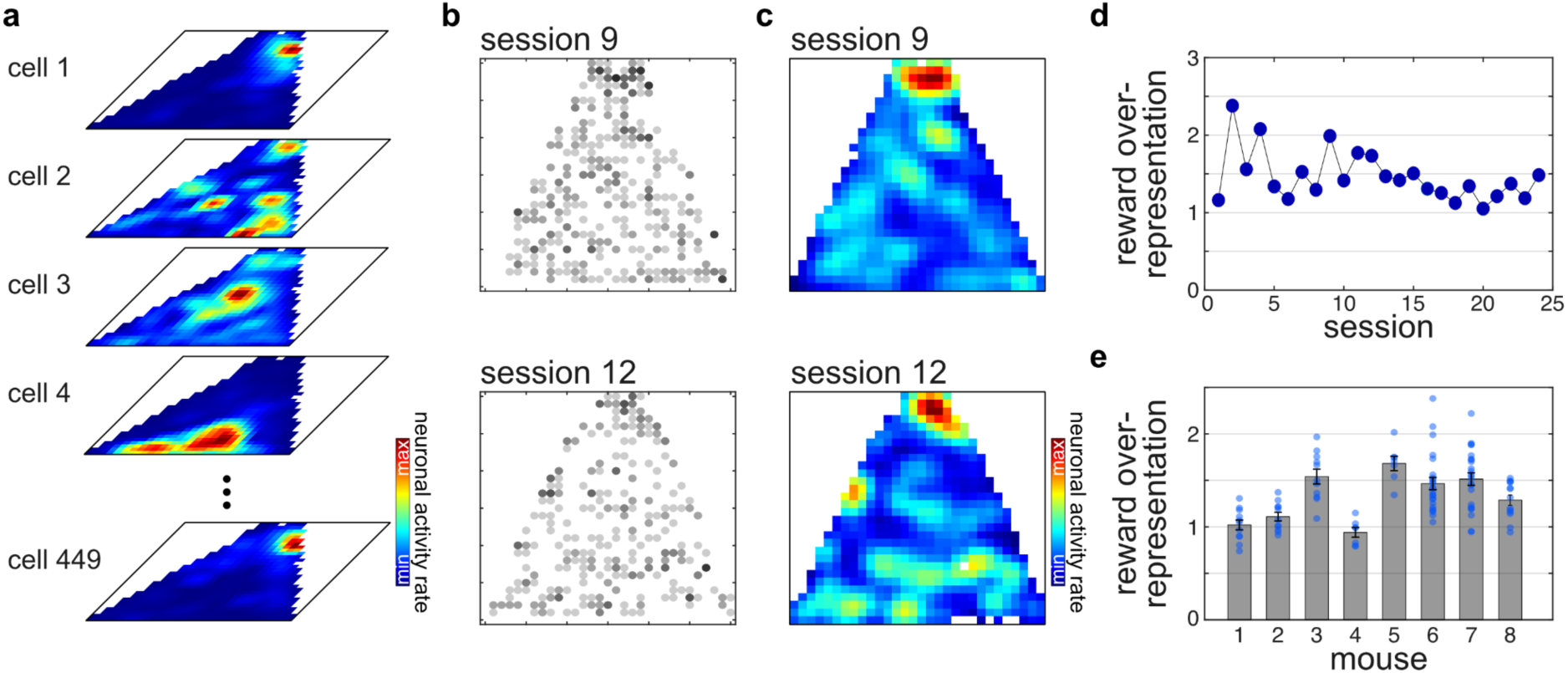
Reward port spatial over-representation. **a,** Sample spatial rate maps for 5 representative cells. **b,** Scatter plot of peak rate maps. Each point represents the position of the peak activity of spatial rate map for each cell. **c,** Density plot of the scatter plots shown in b. **d,** Reward over representation score for all the sessions of mouse #6 are calculated. The score is measured based on the overall representation shown in panel c. It is defined as the average of 10% spatial bins closest to reward normalized by the average of all bins. **e,** Reward over-representation across all mice.

**Extended Data Fig. 7.**
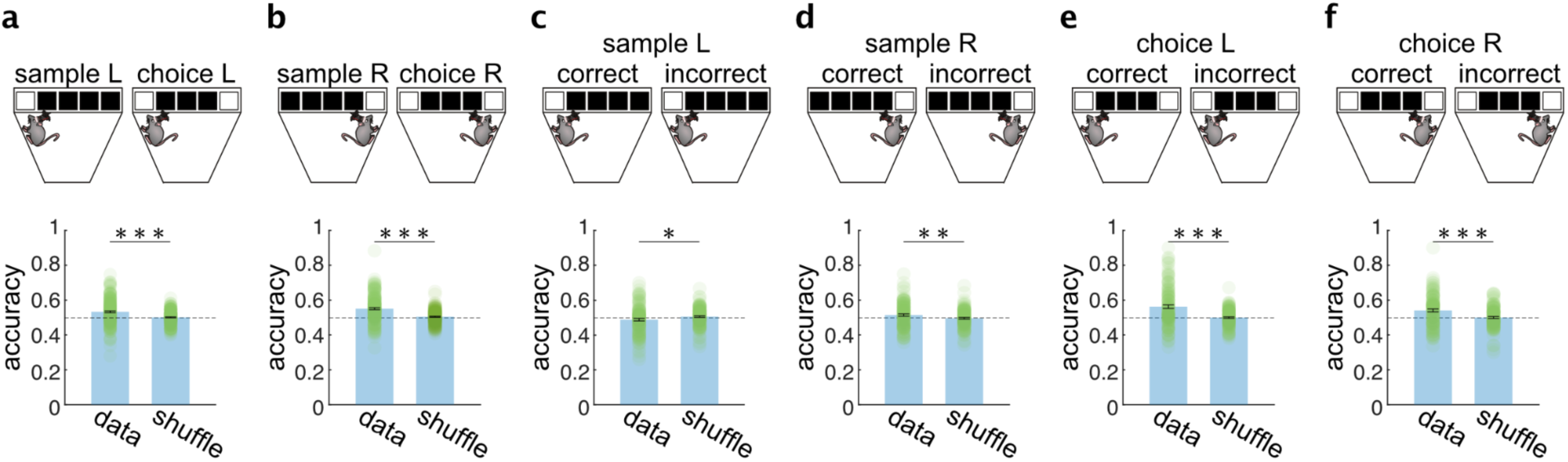
Decoding task-related contextual features while accounting for spatial context. **a,** Task phase decoding (sample vs choice) during left screen contact. Wilcoxon signed-rank test P-value = 9.4979e-10. **b,** Task phase decoding (sample vs choice) during right screen contact. Wilcoxon signed-rank test P-value = 4.3287e-13. **c,** Decoding correctness during left screen sample contact. Wilcoxon signed-rank test P-value = 0.0220. **d,** Decoding correctness during right screen sample contact. Wilcoxon signed-rank test P-value = 0.0083. **e,** Decoding correctness during left screen choice contact. Wilcoxon signed-rank test P-value = 1.1285e-07. **f,** Decoding correctness during right screen choice contact. Wilcoxon signed-rank test P-value = 3.2030e-06.

**Extended Data Fig. 8.**
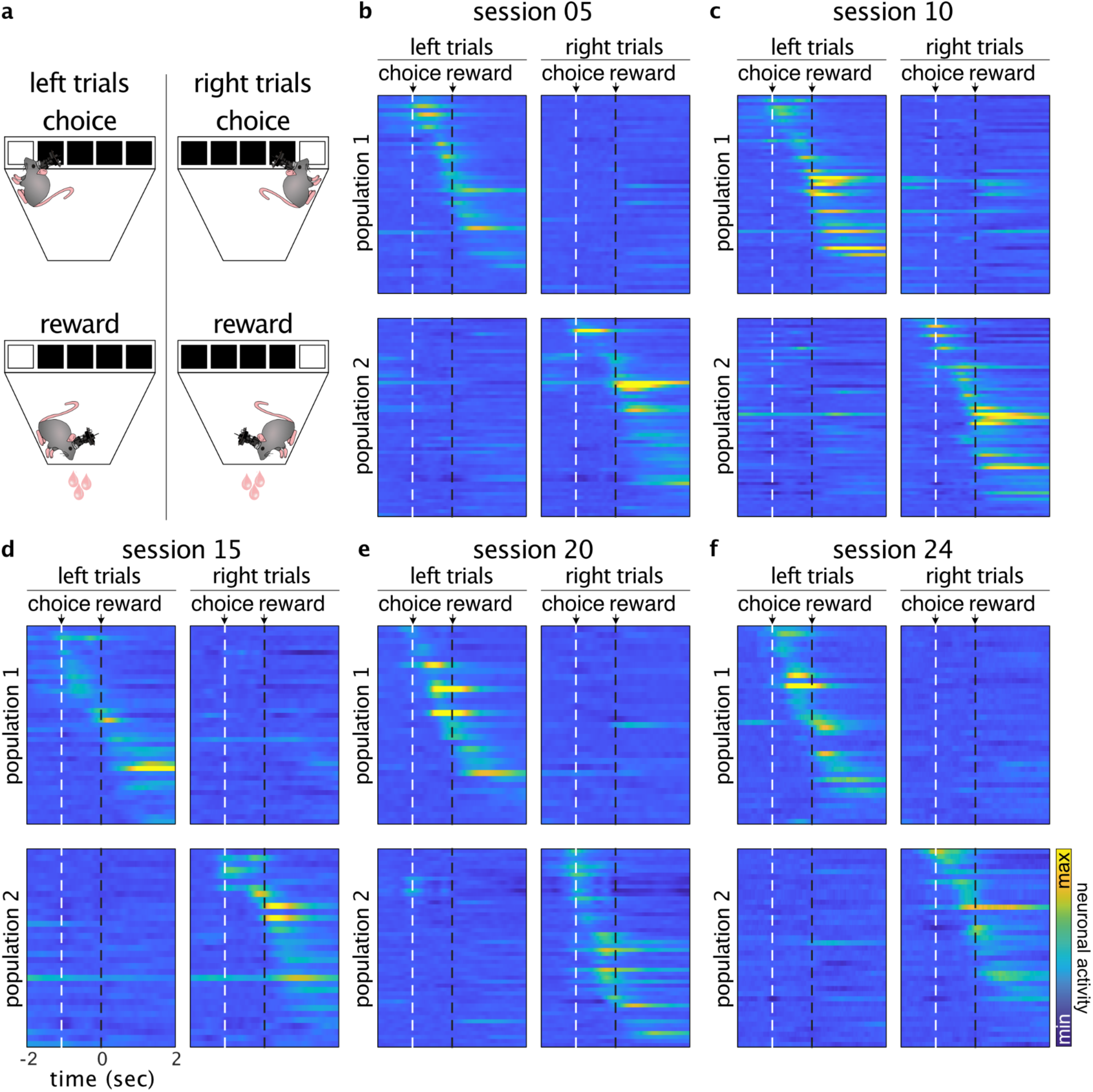
Distinct neuronal sequences cover the choice to reward interval for left and right trials. **a**. A schematic depicting the different trial types and the phases of the task (choice and reward). **b-f**. For each session, the three cell types of interest—screen cells, reward approach cells, and reward cells—are concatenated. Neurons are divided into two groups, Population 1 and Population 2, based on their responses to each trial type. Neurons with higher peak activity during left trials are assigned to Population 1, while those with higher peak activity during right trials are assigned to Population 2. Within each population, cells are sorted by the timing of their peak activity. The white dashed line indicates the choice time, and the black dashed line marks the onset of reward. The time between choice and reward is scaled to one.

**Extended Data Fig. 9.**
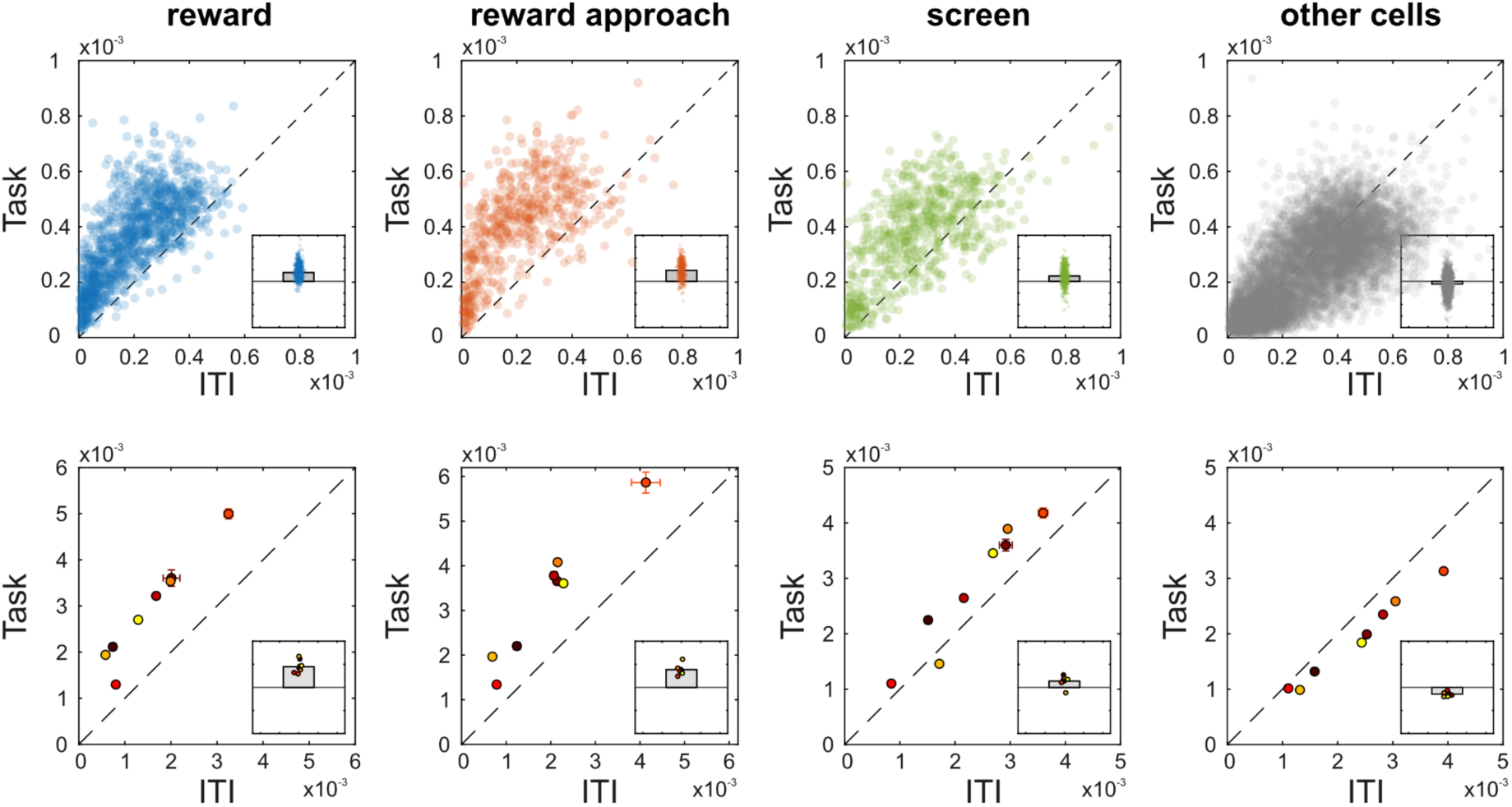
Task vs ITI neuronal responses for different cell types. The neuronal activity (averaged deconvolved calcium traces) for each of the cell types during the task vs ITI is plotted. For all of the cell types, there is a significant difference between task and ITI moments, demonstrating the task-related engagement of the cells. This is not observed for other cells (cells that are not identified as one of the three cell types). Top row: each point is one cell and we have combined all sessions of one mouse (mouse #6). Bottom row: averaged across all cells for each mouse; each point represents one of the mice.

**Extended Data Fig. 10.**
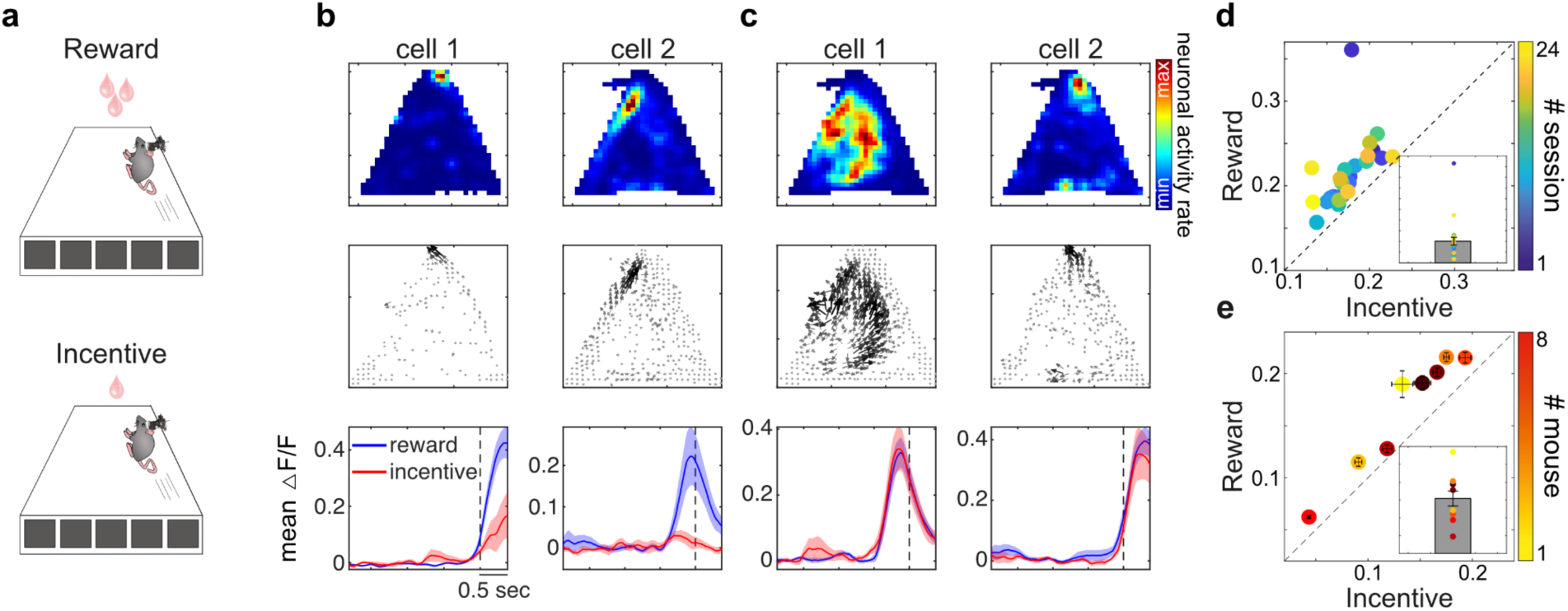
Reward magnitude modulates reward approach neuronal response amplitude. **a,** Schematic of mouse running towards reward (bigger reward) versus incentive (smaller reward). **b,** Two example neurons showing stronger response when mouse is running towards reward (blue) vs incentive (red). **c,** Two example neurons showing similar responses when the mouse is running towards reward (blue) vs incentive (reward). **d,** The average calcium activity in reward approach neurons is higher when the mouse (mouse #6) is approaching a reward compared to incentive. Each point represents a recording session. **e,** similar analysis as d across mice.

**Extended Data Fig. 11.**
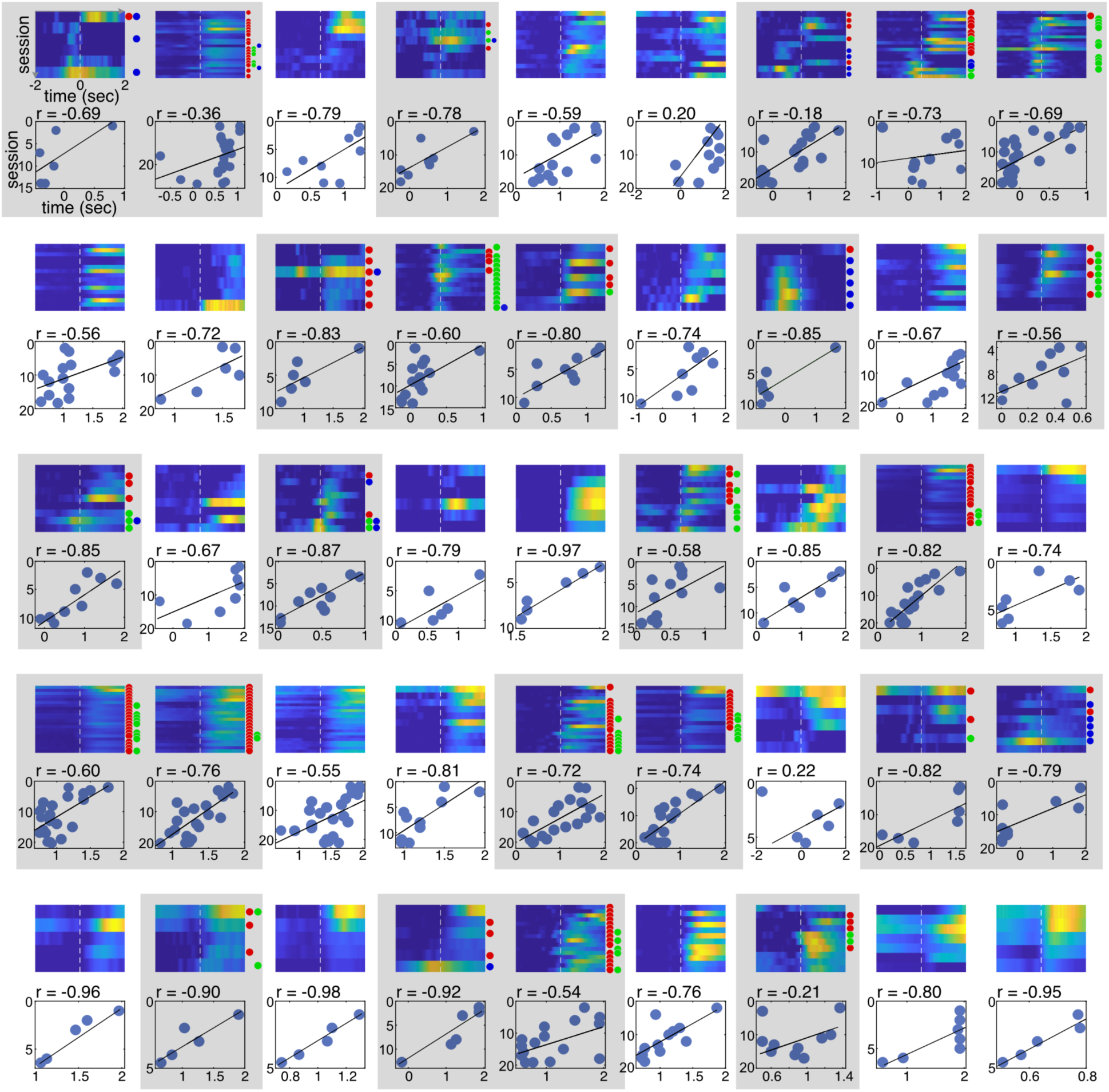
Backward-shifting reward cells. All 45 cells that were identified as backward-shifting reward cells are plotted. Each row represents one session and shows the average of raw calcium traces across all correct trials within that session. 25 of these reward cells are identified as screen or reward approach cells in later days (in gray-shaded box). For the cells in gray shades, the dots in front of each session determine the type of cell in each session; red: reward cell, green: reward approach cell, and blue: screen cell. In some cases the cell shows an extended activity and becomes classified as more than one cell type. Dashed line indicates the onset of the reward. The number above each scatter plot shows the correlation between session number and the timing of reward cell peak activity. The black line is the fitted linear line.

## Notes

### Competing Interest Statement

The authors have declared no competing interest.

### Summary of Updates

We only fixed a couple of typos that existed in the first submission. There is no major change.

